# Differential Metabolic Sensitivity of Insulin-like-response- and mTORC1-Dependent Overgrowth in *Drosophila* Fat Cells

**DOI:** 10.1101/606699

**Authors:** Maelle Devilliers, Damien Garrido, Mickael Poidevin, Thomas Rubin, Arnaud Le Rouzic, Jacques Montagne

## Abstract

The glycolytic/lipogenic axis promotes the synthesis of energetic molecules and building blocks necessary to support cell growth, although the absolute requirement of this metabolic axis must be deeply investigated. Here, we used *Drosophila* genetics and focus on the mTOR signaling network that controls cell growth and homeostasis. mTOR is present in two distinct complexes, mTORC1 and mTORC2. The former directly responds to amino acids and energetic levels, whereas the latter is required to sustain the signaling response downstream of insulin-like-peptide (Ilp) stimulation. Either signaling branch can be independently modulated in most *Drosophila* tissues. We confirm this independency in the fat tissue. We show that ubiquitous over-activation of mTORC1 or Ilp signaling affects carbohydrate and lipid metabolism, supporting the use of *Drosophila* as a powerful model to study the link between growth and metabolism. We show that cell-autonomous restriction of glycolysis or lipogenesis in fat cells impedes overgrowth dependent on Ilp-but not mTORC1-signaling. Additionally, ubiquitous deficiency of lipogenesis (*FASN* mutants) results in a drop in mTORC1 but not Ilp signaling, whereas, at the cell-autonomous level, lipogenesis deficiency affects none of these signals in fat cells. These findings thus, reveal differential metabolic sensitivity of mTORC1- and Ilp-dependent overgrowth. Furthermore, they suggest that local metabolic defects may elicit compensatory pathways between neighboring cells, whereas enzyme knockdown in the whole organism results in animal death. Importantly, our study weakens the use of single inhibitors to fight mTOR-related diseases and strengthens the use of drug combination and selective tissue-targeting.

## INTRODUCTION

Growth of a multicellular organism is coordinated by signaling pathways that adjust intracellular processes to environmental changes. These signaling pathways include the mTOR (mechanistic Target Of Rapamycin) regulatory network that integrates the growth factor response as well as the nutritional and energetic status (Laplante and Sabatini 2012; Howell *et al*. 2013; Lamming and Sabatini 2013; Shimobayashi and Hall 2014; Caron *et al*. 2015; Saxton and Sabatini 2017; Mossmann *et al*. 2018). Activation of this network promotes basal cellular functions, thereby providing building blocks to sustain cellular growth. However, despite a plethora of studies on the mTORC signaling network, the requirement of basal metabolism—glycolytic/lipogenic axis— for cell growth has not been systematically investigated. The *Drosophila* model provides a powerful genetic system to address these issues (Ugur *et al*. 2016), since both the intermediates of this signaling network and the basal metabolic pathways are conserved in the fruit fly (Montagne *et al*. 2001; Hay and Sonenberg 2004; Padmanabha and Baker 2014; Antikainen *et al*. 2017; Wangler *et al*. 2017; Lehmann 2018).

The mTOR protein kinase is present in two distinct complexes, mTORC1 and mTORC2 that comprise raptor and rictor, respectively (Kim *et al*. 2002; Sarbassov *et al*. 2005). Regulation of mTORC1 activity by ATP and amino acids depends on a multi-step process that results in the recruitment of an mTORC1 homodimer at the lysosomal membrane in the vicinity of the small GTPase Rheb (Ras homologue enriched in brain) (Goberdhan *et al*. 2009; Ma and Blenis 2009; Dibble and Manning 2013; Groenewoud and Zwartkruis 2013; Montagne 2016). Rheb stimulates mTORC1 activity (Yang *et al*. 2017), which in turn regulates several downstream targets. S6Kinase1 (S6K1) is one such kinase, sequentially activated through the phosphorylation of its T389 and T229 residues by mTORC1 and by PDK1 (Phosphoinositide-dependent protein kinase 1), respectively (Montagne and Thomas 2004; Magnuson *et al*. 2012). Further, Rheb activation of mTORC1 is repressed by the tumor suppressor TSC (Tuberous sclerosis complex) that comprises subunits TSC1 and TSC2 (Radimerski *et al*. 2002a; Garami *et al*. 2003; Inoki *et al*. 2003a; Dibble *et al*. 2012). The integrity of mTORC2 is required to sustain the downstream insulin-signaling response (Sarbassov *et al*. 2005). Binding of insulin or related peptides (Ilps) to their cognate receptors results in recruitment of class I PI3K (Phosphoinositide 3-kinase) to the membrane. PI3K phosphorylates inositol lipids producing phosphatidylinositol-3,4,5-triphosphate (PIP3) (Engelman *et al*. 2006; Haeusler *et al*. 2018), while the tumor suppressor PTEN acts as a lipid phosphatase to counteract this process (Cully *et al*. 2006; Goberdhan *et al*. 2009). PIP3 constitutes a membrane docking site for the protein kinase Akt whose activity requires the subsequent phosphorylation of its S473 and T308 residues by mTORC2 and PDK1, respectively (Lien *et al*. 2017).

Constitutive activation of mTORC1 in MEFs (Mouse embryonic fibroblasts) has been shown to stimulate a metabolic network, including glycolysis, the pentose phosphate pathway and the biosynthesis of fatty acid (FA) and cholesterol (Duvel *et al*. 2010). Most of the genes encoding glycolytic enzymes are over-expressed in these cells as are those encoding LDH (lactate dehydrogenase) and Pdk1 (Pyruvate dehydrogenase kinase 1; an inhibitor of mitochondrial pyruvate processing). This suggests that mTORC1-activated MEFs potentiate anaerobic glycolysis and repress the tricarboxylic acid (TCA) cycle. Conversely, adipose-specific knockout of raptor to impede mTORC1 formation, results in enhanced uncoupling of mitochondrial activity (Polak *et al*. 2008). The increased lipogenesis observed in mTORC1 stimulated cells depends on a downstream transcriptional regulatory axis involving the cofactor Lipin 1 along with a SREBP (Sterol responsive element binding-protein) family member, which activates genes encoding lipogenic enzymes (Duvel *et al*. 2010; Peterson *et al*. 2011). Congruently, another study revealed that TSC2 mutant cells become addicted to glucose as a result of mTORC1 hyper-activity (Inoki *et al*. 2003b). In addition, inhibition of mTORC1 activity revealed that these TSC2 mutant cells become also dependent on glutamine catabolism (Choo *et al*. 2010); mTORC1 potentiates this catabolism to feed TCA anaplerosis, through 1) a S6K/eIF4B/Myc axis that increases glutaminase protein levels (Csibi *et al*. 2014) and 2) the repression of SIRT4, a mitochondrial sirtuin that inhibits glutamine dehydrogenase (Csibi *et al*. 2013). Besides mTORC1 mediated regulation, Ilp-signaling also impinges on basal metabolism. Intracellular activation of Akt increases ATP levels (Hahn-Windgassen *et al*. 2005; Robey and Hay 2009) through the stimulation of GLUT4-mediated glucose uptake (Jaldin-Fincati *et al*. 2017) and the enhancement of the expression and activity of glycolytic enzymes (Gottlob *et al*. 2001; Houddane *et al*. 2017). Akt also dampens glucose production by suppressing PEPCK (gluconeogenesis), glucose-6-phosphatase (glycogenolyse) and the glycogen synthesis repressor GSK3 (Nakae *et al*. 2001; McManus *et al*. 2005). However, in contrast to mTORC1, Akt also promotes mitochondrial metabolism and oxidative phosphorylations (Gottlob *et al*. 2001; Majewski *et al*. 2004). Conversely, hepatic knockout of the mTORC2 specific-subunit rictor results in constitutive gluconeogenesis and impaired glycolysis and lipogenesis (Hagiwara *et al*. 2012; Yuan *et al*. 2012). Taken together, these studies strongly emphasize the role of mTOR in metabolic-related diseases and in adjusting metabolism to the nutritional and energetic status (Mossmann *et al*. 2018).

In the present study, we investigated the requirement of the glycolytic/lipogenic axis for the cellular growth induced by hyper-activation of mTORC1 signaling and Ilp response in *Drosophila*. As previously demonstrated, mTORC1 and Ilp signaling reside on independent branches in most *Drosophila* tissues (Radimerski *et al*. 2002a; Radimerski *et al*. 2002b; Dong and Pan 2004; Montagne *et al*. 2010; Pallares-Cartes *et al*. 2012). Here, we confirmed this independency in the *Drosophila* fat body (FB), the organ that fulfils hepatic and adipose functions to control body homeostasis (Padmanabha and Baker 2014; Antikainen *et al*. 2017; Lehmann 2018). We show that ubiquitous over-activation of mTOR or Ilp signaling provokes an apparent enhancement of metabolite consumption. Furthermore, our study reveals that metabolic restriction at the organismal level has dramatic consequences on animal survival, but minor effect at the cell-autonomous level, suggesting that within an organism, alternative pathways may operate to compensate local metabolic defects. Nonetheless, at the cell-autonomous level, metabolic restriction can partially restrain overgrowth dependent on hyper-activation of Ilp-but not mTORC1-signaling, indicating that the potential compensatory metabolic pathways do not fully operate in the context of Ilp-signaling stimulation.

## MATERIAL & METHODS

### Genetics and fly handling

Fly strains: *P[w[*+*mC]=tubP-GAL80]LL10,P[ry[*+*t7*.*2]=neoFRT]40A, daughterless(da)-gal4, tub-gal80*^*ts*^, *UAS-Dcr-2* (Bloomington Stock Center); *FASN*^*1-2*^ (Garrido *et al*. 2015); *mTOR*^*ΔP*^ (Zhang *et al*. 2000); *mTOR*^*2L1*^ and *PTEN* (Oldham *et al*. 2000); *EP(UAS)-Rheb* (Stocker *et al*. 2003);); inducible interfering RNA (*UAS-RNAi*) lines to *PTEN* (NIG 5671R-2), *FASN1* (VDRC 29349), *PFK1* (VDRC 3017), *PK* (VDRC 49533) *PDH* (VDRC 40410), *LDH* (VDRC 31192) (Dietzl *et al*. 2007). The *Minute* stock used was previously referred to as *FRT40/P(arm-LacZ w*^+^) (Bohni *et al*. 1999) but exhibit both developmental delay and short and slender bristles, typically reported as *Minute* phenotype (Morata and Ripoll 1975). To generate MARCM clones in the *Minute* background, these flies were recombined with the *P[w[*+*mC]=tubP-GAL80]LL10,P[ry[*+*t7*.*2]=neoFRT]40A* chromosome.

The standard media used in this study contained agar (1g), polenta (6g) and yeast (4g) for 100ml. Lipid-(beySD) and sugar-complemented media were prepared as previously described (Garrido *et al*. 2015).

To select *FASN*^*1-2*^ mutant larvae, we used a GFP-labelled CyO balancer chromosome. Flies were let to lay eggs on grape juice plates for less than 24 hrs. Then, some beySD media was put in the middle of the plates; larvae that do not express GFP were collected the next day and transferred to fresh tubes. Prepupae were collected once a day to evaluate developmental delay and to measure body weight.

### Molecular biology and Biochemistry

To test RNAi-knockdown efficacy to the glycolytic enzymes (Figure S2), *UAS-Dcr-2;da-gal4,tub-gal80*^*ts*^ virgin females were mated with UAS-RNAi males. Flies were let to lay eggs overnight and tubes were kept at 19°C for two days. Tubes were then transferred at 29°C and two days later, larvae of roughly the same size were collected. Reverse transcription and quantitative PCR were performed as previously described (Parvy *et al*. 2012).

Protein extracts for western-blotting were prepared as previously described (Montagne *et al*. 2010). Antibody used in for western-blotting have been previously described (Montagne *et al*. 2010) or commercially provided for Akt (Cell signaling 4054).

For metabolic measurements, parental flies were let to lay eggs in tubes for less than 24 hrs at 25°C. Tubes were then transferred at 29°C to strengthen the gal4/UAS effect, and using a *UAS-Dcr-2* to strengthen the RNAi effect. Larvae were either maintained in the same tubes or selected prior to L2/L3 transition and transferred on 20%-SSD. Collection of prepupae and metabolic measurements were performed as previously described (Garrido *et al*. 2015).

### Clonal analysis

All the clones were generated using the MARCM strategy (Lee and Luo 2001). Parental flies were let to lay eggs at 25°C for seven hrs. Tubes were then heat shocked for 65 minutes in a water bath at 38°C so that recombination happens while FB precursor cells are in dividing process. FB from feeding larvae at the end of the L3 stage where dissected, fixed, membranes were labelled with phalloidin and nuclei with DAPI, and FB were mounted as previously described (Garrido *et al*. 2015). Image acquisitions were obtained using a Leica SP8 confocal laser-scanning microscope. For immuno staining the phospho-S6 antibody has been previously described (Romero-Pozuelo *et al*. 2017) and the phospho-Akt commercially provided (Cell signaling 4054). The cell size calculation have been performed as previously described (Garrido *et al*. 2015) and correspond to a set of experiments that spanned a two-year period. It represent too many replicates, so that it was not possible to make them at the same time. Therefore, for the graphs of cell size measurement (Figure 1M, 5M and 7M), values are reused when they correspond to the same genotype and conditions. This allows a direct comparison between the experiments.

**Figure 1:**
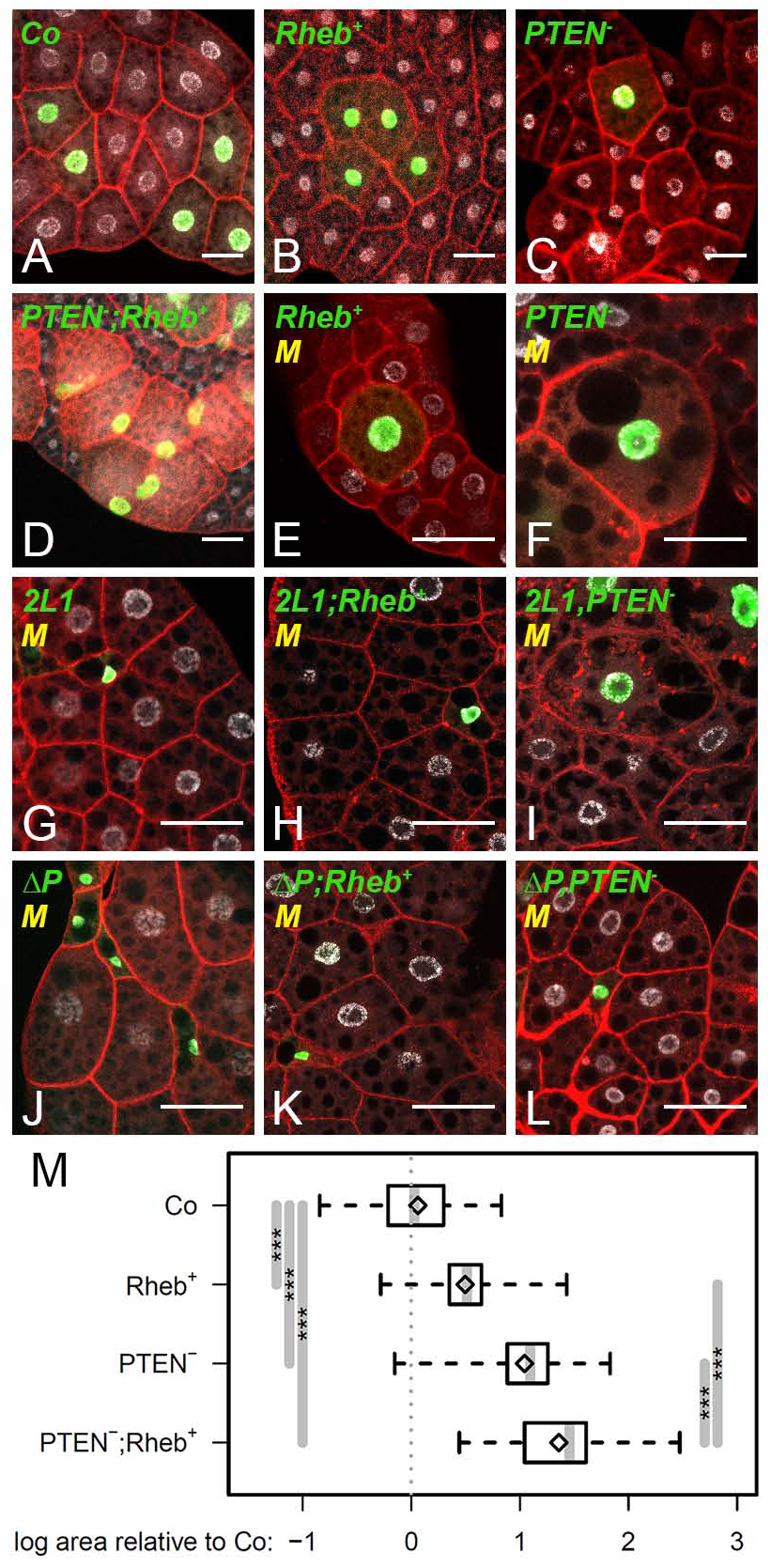
mTORC1- and Ilp-dependent growth in FB cells. (**A-L**) MARCM clones labeled by GFP (green) in the FB of L3 larvae. Nuclei were labeled with DAPI (silver) and membranes with phalloidin (red). Control (A), *Rheb*^+^ (B), *PTEN*^−*/*−^ (C) and *PTEN*^−*/*−^;*Rheb*^+^ (D) clones were generated in a wild type background. *Rheb*^+^ (E), *PTEN*^−*/*−^ (F), *mTOR*^*2L1*^ (G) *mTOR*^*2L1*^, *Rheb*^+^ (H) *mTOR*^*2L1*^,*PTEN*^−*/*−^ (I), *mTOR*^*ΔP*^ (J), *mTOR*^*ΔP*^,*Rheb*^+^ (K) and *mTOR*^*ΔP*^,*PTEN*^−*/*−^ (L) clones were generated in a *Minute* (*M*) background. Scale bars: 50μm. (**M**) Relative size of control (Co), *Rheb*^+^, *PTEN*^−*/*−^, and *PTEN*^−*/*−^;*Rheb*^+^ clonal cells generated in a wild type background.

### Statistical analysis

Statistical analyses were performed with R version 3.4.4, scripts are available on request. Significance for the statistical tests was coded in the following way based on the p-values: ***: 0 < p < 0.001; **: 0.001 < p < 0.01; *: 0.01 < p < 0.05. P-values were corrected for multiple testing by a Holm-Bonferroni method (Holm 1079). Clone sizes were analyzed with a mixed-effect linear model on the logarithm of cell area, considering the treatment (Genotype and Sucrose conditions) as a fixed effect and Series/Larva as random effects (Figures 1, 5, and 7, Table S1). The reported effects (and the corresponding P-values) were obtained from the difference between the (log) area of marked clonal cells and that of control surrounding cells from the same treatment, by setting the appropriate contrast with the “multcomp” package (Hothorn *et al*. 2008), according to the pattern: EA,B = log(MA) – log(WA) – [log(MB) – log(WB)], where EA,B is the difference between treatments (genotype and sucrose levels) A and B, MA and MB standing for the area of marked cells, and WA, WB for the area of control cells in those treatments. This is equivalent to testing whether marked/control cell area ratios differ between treatments. PS6+ clone frequencies were treated as binomial measurements in a mixed-effect generalized linear model “lme4” package (Bates *et al*. 2015), featuring Genotype as a fixed effect, and Series/Larva as random effects. Both datasets of pupal weights were analyzed independently with linear models including Sex, Genotype, and Sucrose level effects and all their interaction terms (Figure 3A-B and Table S3 for PTEN knockdown and Rheb overexpression; Figure 6B and Table S4 for *FASN*^*1-2*^ mutants). TAG, Protein, Glycogen, and Threalose concentrations were also analyzed with linear models involving Genotype, Sucrose level, and their interactions as fixed effects (Figure 3 and Table S3).

**Figure 2:**
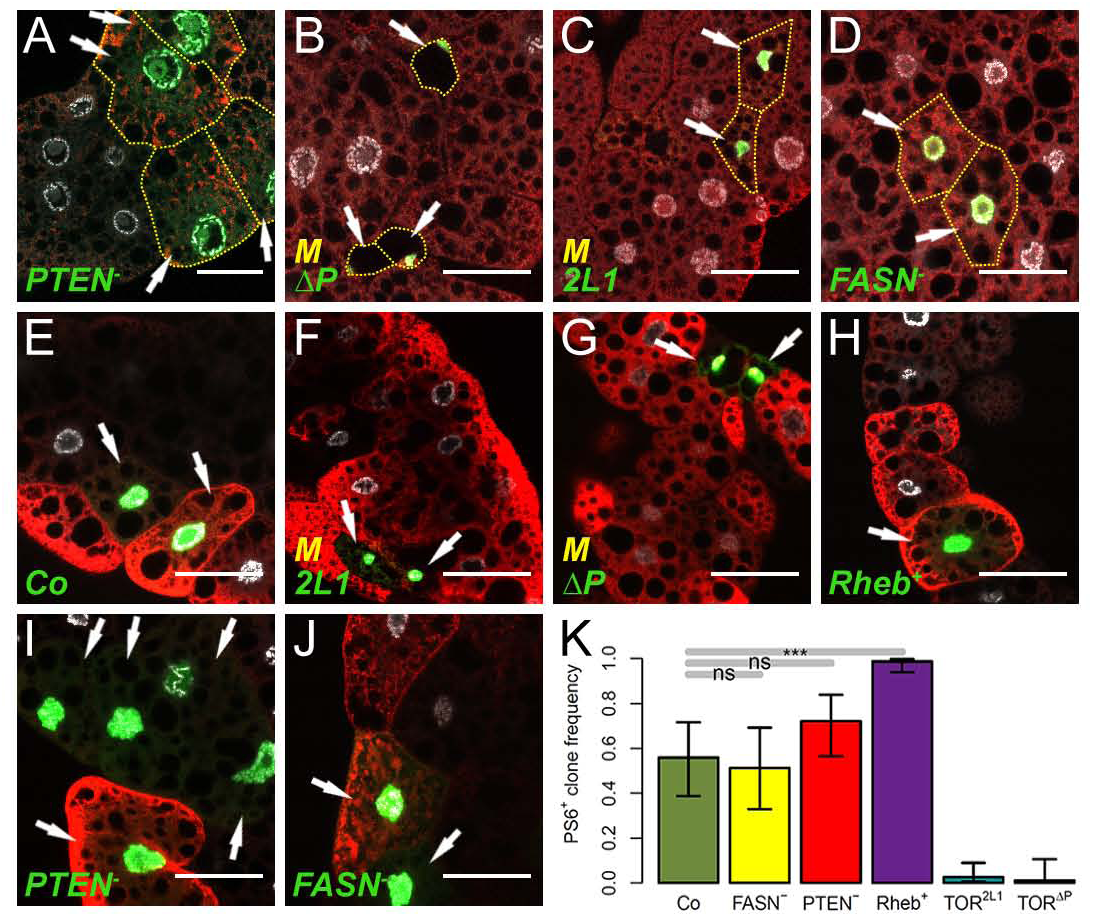
mTORC1 and Ilp signaling activity in FB cells. (**A-J**) MARCM clones labeled by GFP (green) in the FB of L3 larvae. Clones were generated in a wild type (A,D,E,H,I,J) or a *Minute* (B,C,F,G) background and nuclei were labeled with DAPI (silver). FB tissues with *PTEN*^−*/*−^ (A), *mTOR*^*ΔP*^ (B), *mTOR*^*2L1*^ (C) and *FASN*^*1-2*^ (D) clones were stained with a phospho-AKT antibody. FB tissues with control (E), *mTOR*^*2L1*^ (F), *mTOR*^*ΔP*^ (G) *Rheb*^+^ (H), *PTEN*^−*/*−^ (I) and *FASN*^*1-2*^ (J) clones were stained with a phospho-S6 antibody. Scale bars: 50μm. (**K**) Percentage of P-S6 positive clones with respect to the total number of MARCM clones for control, *FASN*^*1-2*^, *PTEN*^−*/*−^, *Rheb*^+^, *mTOR*^*2L1*^ and *mTOR*^*ΔP*^ genotypes.

**Figure 3:**
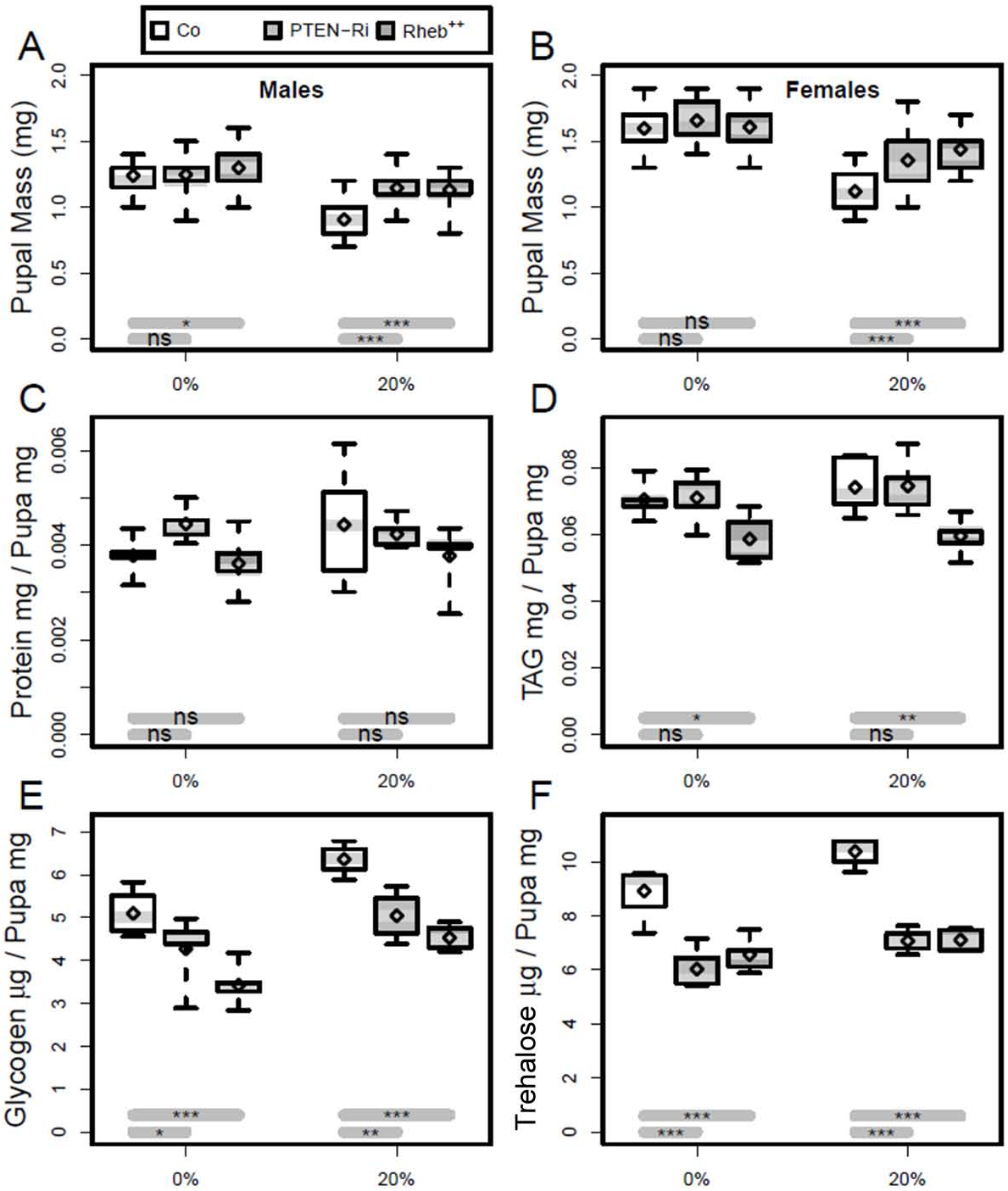
Enhanced mTORC1 or Ilp signaling affects larval metabolism. (**A-B**) Body weight of female (A) and male (B) prepupae formed from larvae fed either a standard (0%) or a 20%-SSD (20%) as from the L2/L3 transition. (**C-F**) Measurement of total protein (C), TAG (D), glycogen (E) and trehalose (F) levels in prepupae fed either a standard or a 20%-SSD. Prepupae used in these measurements were the F1 progeny from *da-gal4* virgin females mated to either control (Co), *EP(UAS)-Rheb* (*Rheb*^++^) or *UAS-PTEN-RNAi* (*PTEN-Ri*) males.

**Figure 4:**
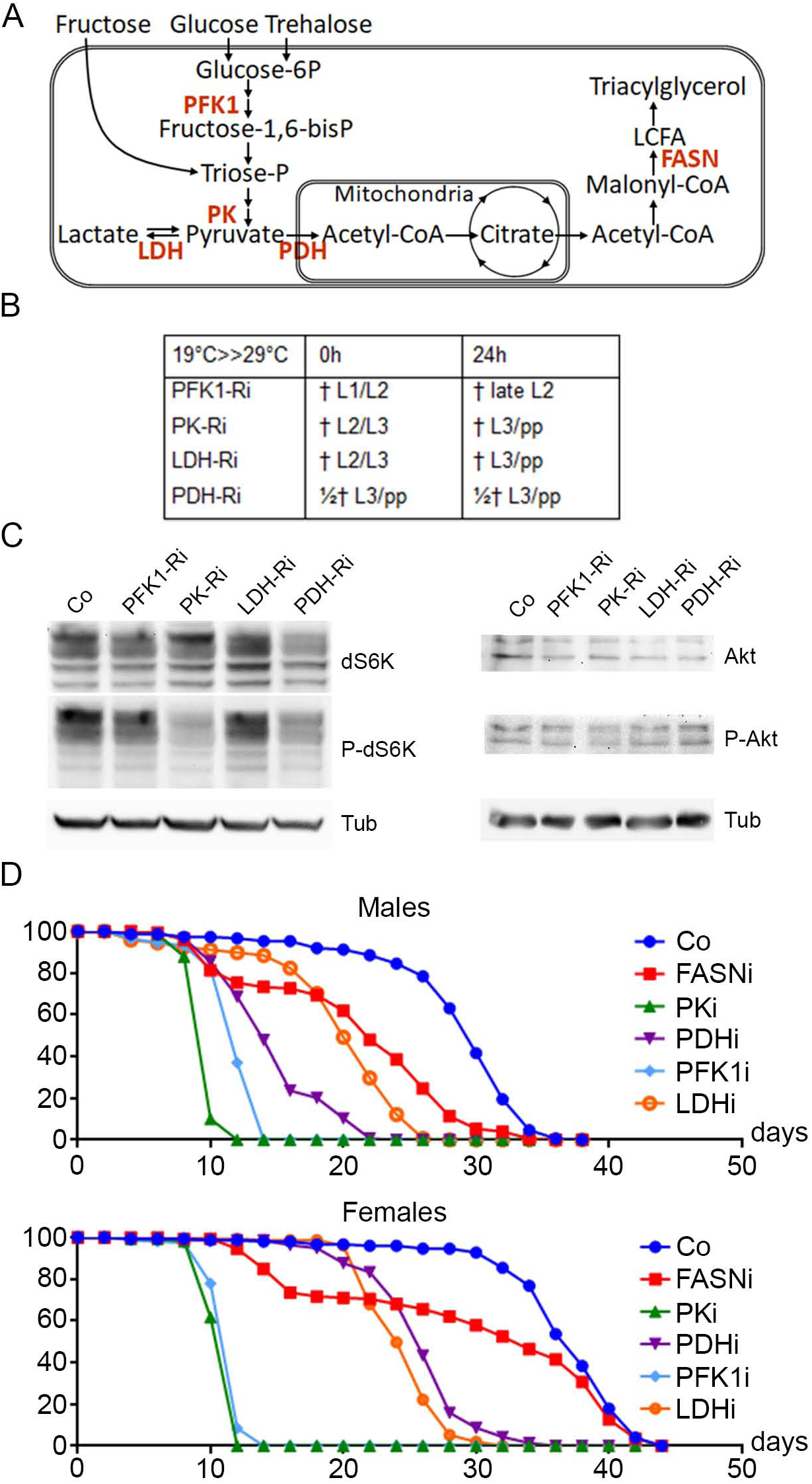
Glycolysis knockdown in whole organisms. (**A**) Scheme of basal metabolism. Glucose and trehalose enter glycolysis as glucose-6P, whereas fructose follows a distinct pathway to triose-P. Enzymes investigated in the present study are marked in red. (**B**) Phenotype of ubiquitous RNAi knockdown of PFK1, PK, LDH and PDH. Flies were left to lay eggs overnight either at 29°C (column 0h) or at 19°C and transferred to 29°C the day after (column 24h); then development proceeded at 29°C (i.e. the temperature that inactivates Gal80). (**C**) Western-blot analysis of total (top) or phosphorylated (mid) dS6K (left) or Akt (right) proteins; tubulin (bottom) was used as a loading control. Protein extracts were prepared with late L2 control larvae (Co) or L2 larvae expressing RNAi against the indicated metabolic enzymes. (**E-F**) Survival at 29°C of male (top) and female (bottom) control flies or flies expressing RNAi against the indicated metabolic enzymes as from adult eclosion.

**Figure 5:**
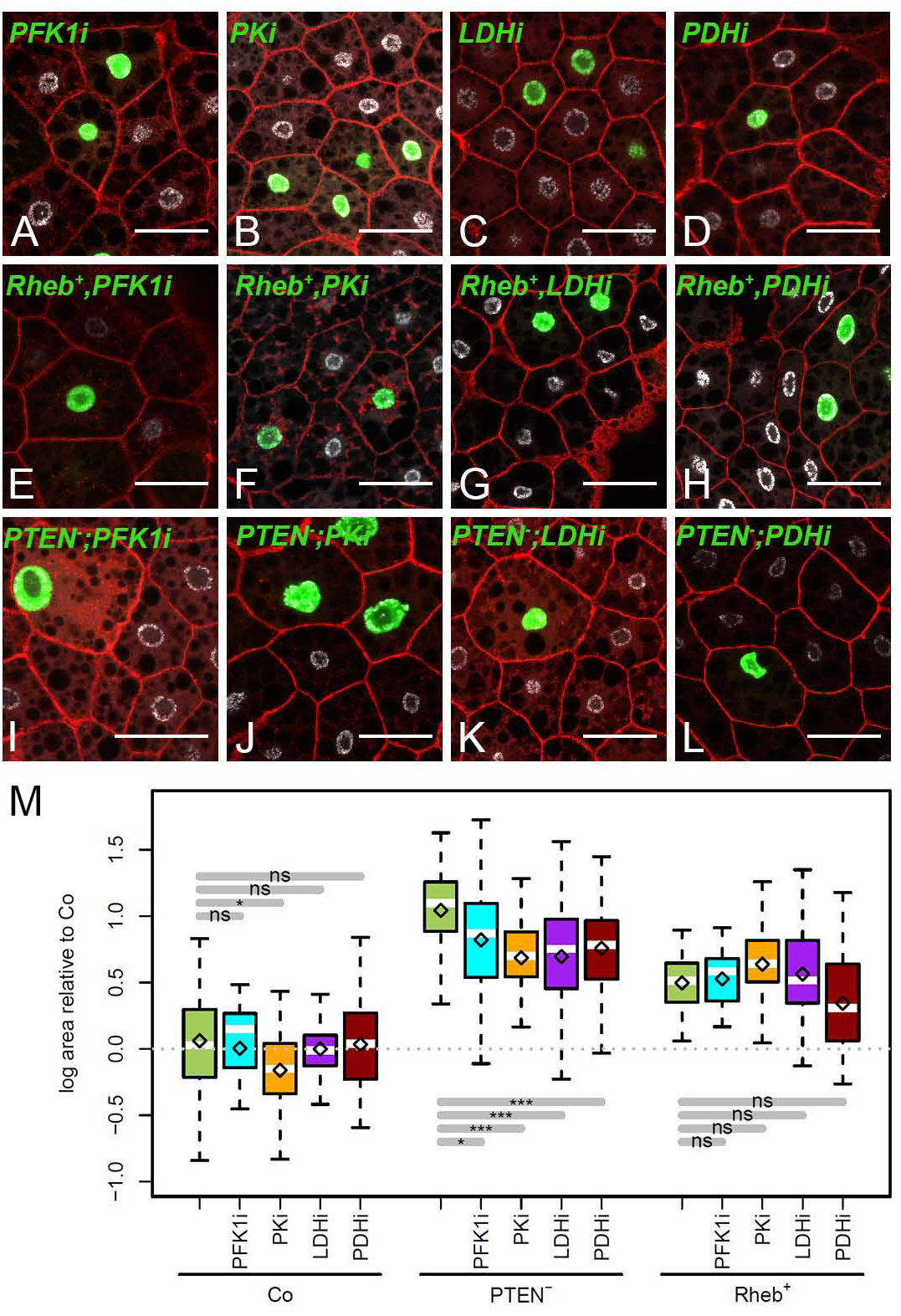
Cell-autonomous requirement of glycolysis for Ilp-but not mTORC1-dependent overgrowth. (**A-G**) MARCM clones labeled by GFP (green) in the FB of L3 larvae. Nuclei were labeled with DAPI (silver) and membranes with phalloidin (red). Genotypes of MARCM clones are: *PFK1-RNAi* (A), *PK-RNAi* (B), *LDH-RNAi* (C), *PDH-RNAi* (D), *Rheb*^+^,*PFK1-RNAi* (E), *Rheb*^+^,*PK-RNAi* (F), *Rheb*^+^,*LDH-RNAi* (G), *Rheb*^+^,*PDH-RNAi* (H), *PTEN*^−*/*−^;*PFK1-RNAi* (I), *PTEN*^−*/*−^;*PK-RNAi* (J), *PTEN*^−*/*−^;*PDH-RNAi* (K) and *PTEN*^−*/*−^;*PDH-RNAi* (L). Scale bars: 50μm. (**M**) Relative size of clonal cells corresponding to the clones shown in A-L, and in Figure 1A for control (Co).

**Figure 6:**
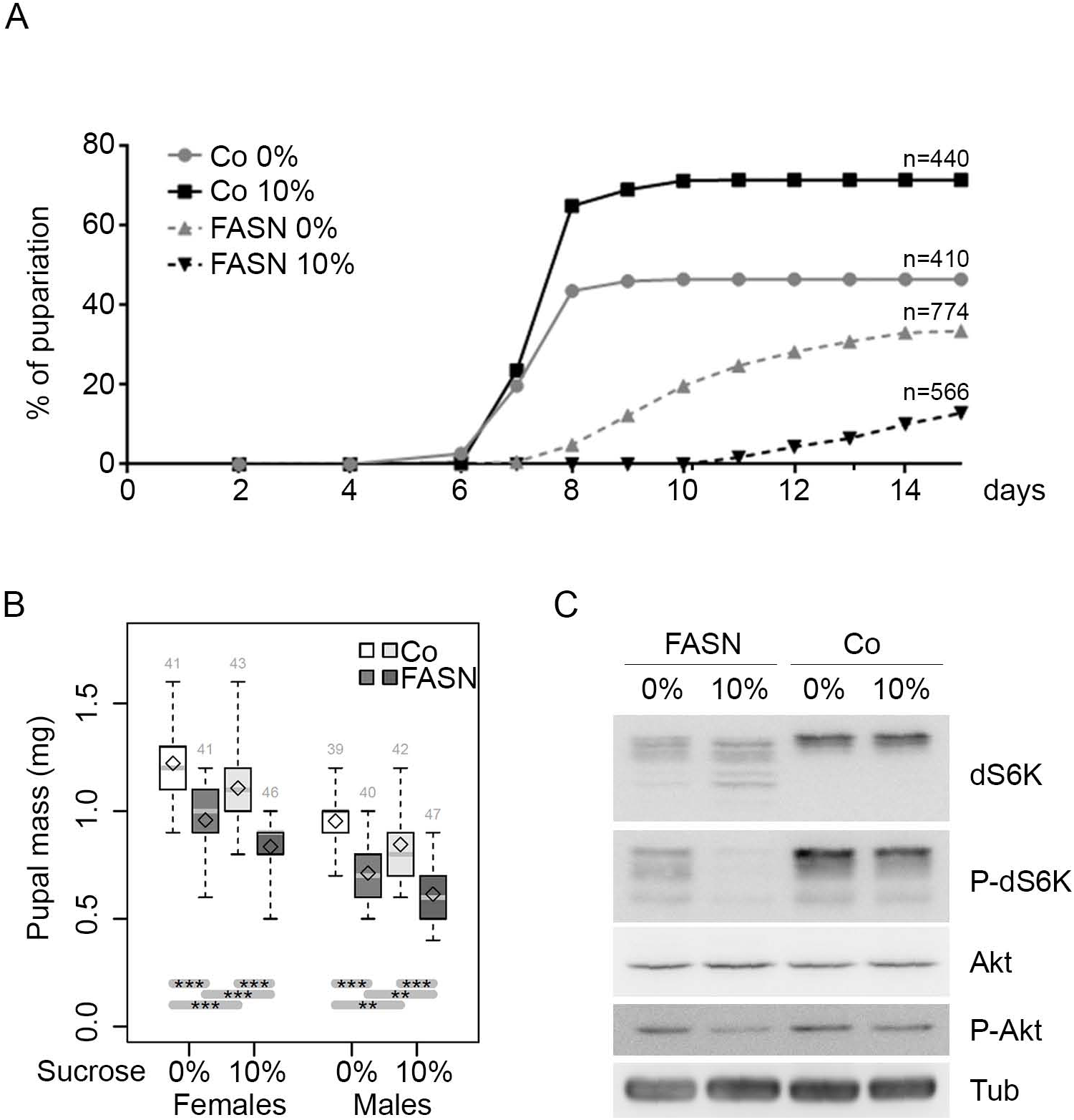
*FASN*^*1-2*^ mutation affects developmental growth and mTORC1 signaling. (**A**) Developmental duration from egg laying to metamorphosis onset of *w*^*1118*^ control (Co) and *FASN*^*1-2*^ (FASN) larvae fed either a beySD (0%) or a 10% sucrose-supplemented-beySD as from the L2/L3 transition (10%); n: total number of larvae collected for each condition. (**B**) Prepupal weight of females (left) and males (right) as listed in 6A; the numbers of weighted prepupae are indicated above each sample. (**C**) Western-blot analysis of (from top to bottom) total dS6K, phosphorylated dS6K, total Akt, phosphorylated Akt and total tubulin as a loading control. Protein extracts were prepared from feeding L3 larvae prior to the wandering stage as listed in 6A. For each condition, at least 30 larvae were used to prepare protein extracts.

**Figure 7:**
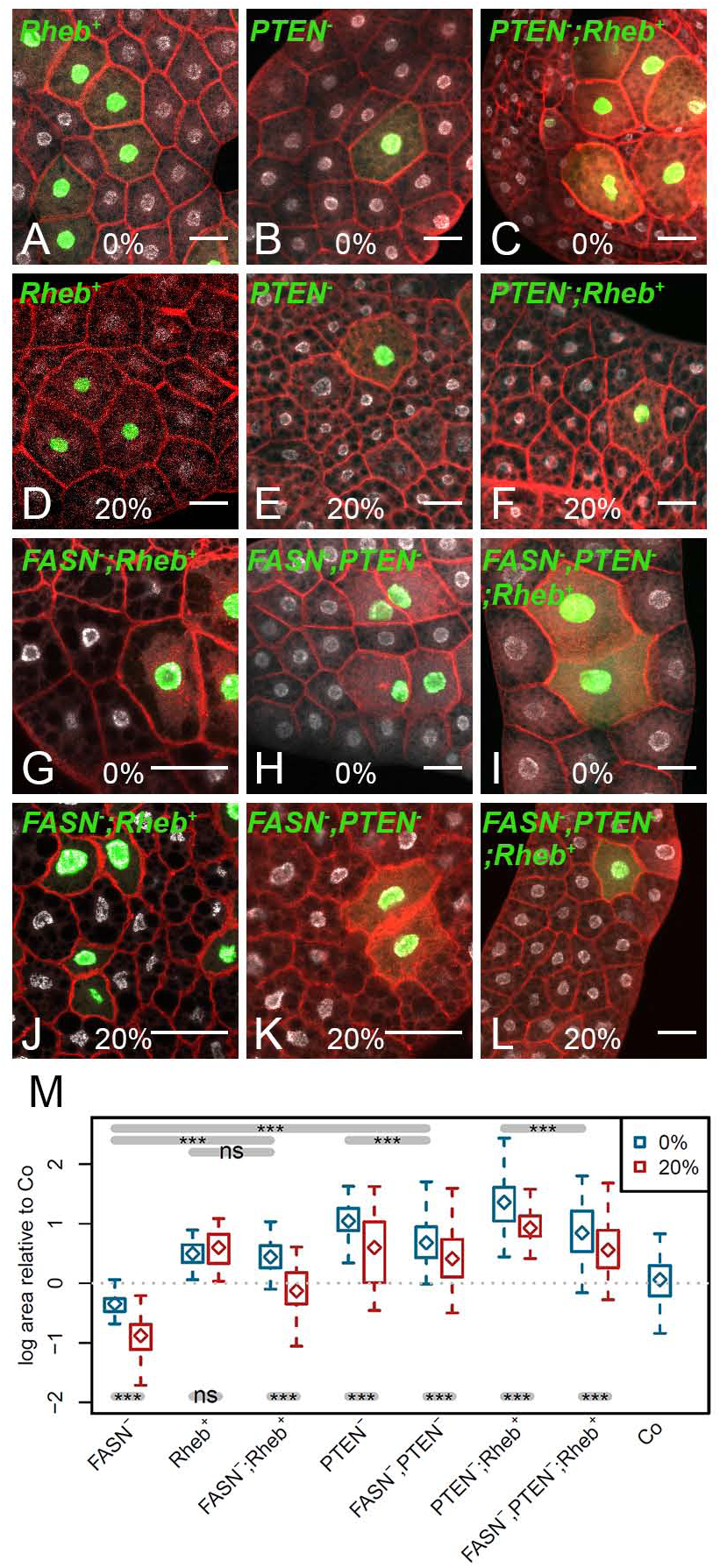
Cell-autonomous requirement of FASN activity for Ilp-but not mTORC1-dependent overgrowth. (**A-L**) MARCM clones labeled by GFP (green) in the FB of L3 larvae fed either a standard (A-C, G-I) or a 20%-SSD (D-F, J-L). Nuclei were labeled with DAPI (silver) and membranes with phalloidin (red). Genotypes of MARCM clones are: *Rheb*^+^ (A,D), *PTEN*^−*/*−^ (B,E) *PTEN*^−*/*−^,*Rheb*^+^ (C,F), *FASN*^*1-2*^;*Rheb*^+^ (G,J) *FASN*^*1-2*^,*PTEN*^−*/*−^ (H,K) and the *FASN*^*1-2*^,*PTEN*^−*/*−^;*Rheb*^+^ (I,L). Scale bars: 50μm. (**M**) Relative size of clonal cells corresponding to the clones shown in A-L and in Figure S1 for *FASN*^*1-2*^ and Figure 1A for control (Co).

### Data and reagent availability statement

Fly stocks are available upon request. Supplementary materials include Figures S1-S2, Tables S1-S4 and supdata/script files available on the GSA figshare portal.

## RESULTS

### mTORC1 and Ilp signaling independency in the fat body

Activating either the mTORC1 or the Ilp signaling branch can be performed by overexpressing Rheb or depleting PTEN, respectively. To investigate this independency in the FB, we generated somatic clones either over-expressing Rheb (*Rheb*^+^) (Stocker *et al*. 2003) or homozygote for a *PTEN* mutation (*PTEN*^−*/*−^) (Oldham *et al*. 2000). The precursors of FB cells divide in the embryo; during larval life, the differentiated cells do not divide but endoreplicate their DNA content to reach a giant size (Edgar and Orr-Weaver 2001). Therefore, to precisely evaluate the effect on cell growth, somatic recombination events were induced during embryogenesis at the stage of proliferation of the FB cell precursors and the resulting MARCM clones were analyzed in the FB of late feeding L3 larvae, prior to the wandering stage that precedes metamorphosis entry. Both *PTEN*^−*/*−^ and *Rheb*^+^ clonal cells were bigger than the surrounding control cells and this cell size effect was dramatically increased in *PTEN*^−*/*−^;*Rheb*^+^ combined clones (Figure 1A-D and 1M). We next analyzed this growth increase in the context of the previously described *mTOR*^*2L1*^ and *mTOR*^*ΔP*^ mutations. However, we could not find mutant clones in the FB. Consistent with previous studies reporting that mTOR is critically required for cell growth of endoreplicative tissues (Oldham *et al*. 2000; Zhang *et al*. 2000), we reasoned that these clonal cells were likely eliminated by cell competition (Morata and Ripoll 1975). Thus, we generated somatic clones in a *Minute* background to slow down the growth of the surrounding control cells. In these conditions, *mTOR* mutant clones could indeed be recovered. Both *mTOR*^*2L1*^ and *mTOR*^*ΔP*^ mutant cells exhibited a dramatic size reduction (Figure 1G and 1J) and this phenotype was dominant in *Rheb*^+^ combined clonal cells (compare Figure 1E to 1H and 1K). In contrast, *mTOR*^*ΔP*^ but not *mTOR*^*2L1*^ exhibited a clear dominant phenotype over the *PTEN*^−*/*−^ mutation; the size of *mTOR*^*ΔP*^,*PTEN*^−*/*−^ clonal cells was dramatically reduced, whereas *mTOR*^*2L1*^,*PTEN*^−*/*−^ clonal cells were giant (compare Figure 1F to 1I and 1L). These findings indicate that the *mTOR*^*2L1*^ mutation affects mTORC1 but not ilp signaling, whereas *mTOR*^*ΔP*^ affects both signaling branches.

Next, we used phospho-specific antibodies in immunostaining assays to analyze the phosphorylation of Akt (P-Akt) and of the dS6K target, ribosomal protein rpS6 (P-S6). In *PTEN*^−*/*−^ clonal cells, we observed an increase in the P-Akt intracellular signal (Figure 2A). Importantly the P-Akt intracellular signal was absent in *mTOR*^*ΔP*^ cells (Figure 2B) but not affected in *mTOR*^*2L1*^ cells (Figure 2C). Staining with the rpS6 phospho-specific antibody revealed a patchy signal, with only a subset of cells expressing the P-S6 signal in the FB (Figure 2E-J), a pattern previously described in the wing imaginal disc (Romero-Pozuelo *et al*. 2017). Therefore, to evaluate mTORC1 activity, we measured the ratio of P-S6 positive cells among the population of GFP^+^ clonal cells. For control clones, only labeled by GFP, about half of them were P-S6 positive (Figure 2E and 2K), whereas most of the *mTOR*^*2L1*^ and *mTOR*^*ΔP*^ clones were P-S6 negative (Figure 2F, 2G and 2K). Importantly, almost all the *Rheb*^+^ cells were P-S6 positive (Figure 2H and 2K), whereas the ratio of P-S6 positive cells was slightly but not significantly increased in the *PTEN*^−*/*−^ cell population (Figure 2I and 2K). Taken together, these findings confirm that mTORC1 and Ilp signaling operate independently in FB cells and reveal that the *mTOR*^*2L1*^ mutation affects only mTORC1, whereas the *mTOR*^*ΔP*^ mutation affects both signaling branches.

### Activating mTORC1 or Ilp signaling impacts basal metabolism

A number of studies support the notion that the mTOR signaling network controls metabolism to sustain cellular growth. To evaluate how mTORC1 and Ilp affect basal metabolism in *Drosophila*, we analyzed various metabolites in whole animals that express the ubiquitous *da-gal4* driver to direct Rheb overexpression (*Rheb*^++^) or PTEN knockdown by RNA interference (*PTEN-RNAi*). Larvae were fed either a standard or a 20%-sucrose supplemented diet (20%-SSD) and 0-5h prepupae were collected, as this is a convenient phase to stage the animals after the feeding period. When fed a standard diet, a high rate of lethality was observed for *Rheb*^++^ and *PTEN-RNAi* larvae, although a sufficient number of prepupae could be collected for metabolic analysis. In contrast, none of the *Rheb*^++^ and *PTEN-RNAi* larvae reached the prepupal stage when fed a 20%-SSD. Nonetheless, when *Rheb*^++^ and *PTEN-RNAi* larvae were fed a standard diet during early larval life and transferred onto a 20%-SSD at the L2/L3 molting transition, we could recover a few prepupae for metabolic measurements. For both males and females fed a standard diet, the body weight of *Rheb*^++^ and *PTEN-RNAi* prepupae was roughly similar to that of controls (Figure 3A and 3B). Conversely, providing a 20%-SSD resulted in a drop of the prepupal weight of control animals that was significantly compensated in *Rheb*^++^ and *PTEN-RNAi* prepupae (Figure 3A and 3B).

Next, we measured the total amounts of protein, triacylglycerol (TAG), glycogen and trehalose—the most abundant circulating sugar in *Drosophila*. Although variations in protein levels were observed, none of them were statistically significant (Figure 3C). TAG levels in control prepupae were not affected by sucrose supplementation and did not vary in *PTEN-RNAi*, but were significantly decreased in *Rheb*^++^ animals (Figure 3D). Feeding larvae a 20%-SSD since the L2/L3 molting transition resulted in a marked in increase in glycogen and trehalose levels in control prepupae (Figure 3E-F). In *Rheb*^++^ and, in lower extent, in *PTEN-RNAi* prepupae, glycogen levels were significantly lower than those measured in controls (Figure 3E). Finally, trehalose levels were strongly decreased in both *Rheb*^++^ and *PTEN-RNAi* prepupae fed either a standard or a 20%-SSD as compared to the control (Figure 3F). Taken together, these findings suggest that a ubiquitous increased activity of either mTORC1 or Ilp signaling provokes an apparent increase in metabolite consumption. This metabolic rate is correlated with a relative increase in body weight for larvae fed a 20%-SSD, but not for those fed a standard diet. We previously observed that increasing dietary sucrose induced a reduction in food intake (Garrido *et al*. 2015) that may account for the body weight reduction of control animals. Potentially, food intake could be less affected in *Rheb*^++^ and *PTEN-RNAi* animals, thereby leading to a compensatory effect on body weight. Measuring food intake in *Rheb*^++^ or *PTEN-RNAi* larvae was not applicable since most of them die during larval stage and thus, terminate feeding earlier. In sum, our data indicates that basal metabolism is altered in the few *Rheb*^++^ or *PTEN-RNAi* larvae that survive and further suggests that in most cases stronger metabolic disruption happened, resulting in lethal homeostatic defects.

### Knocking-down glycolysis at the whole body level

Since manipulating mTOR resulted in a decrease in the levels of TAG and glycogen stores and of circulating trehalose (Figure 3), we asked whether the basal energetic metabolism affected mTORC1- and/or Ilp-signaling. First, we ubiquitously expressed interfering RNA against phosphofructokinase1 (*PFK1-RNAi*), pyruvate kinase (*PK-RNAi*) pyruvate dehydrogenase (*PDH-RNAi*) and lactate dehydrogenase (*LDH-RNAi*). PFK1 catalyzes the third glycolytic reaction to form fructose 1,6-bisphophate; PK catalyzes the final glycolytic reaction to form pyruvate; PDH directs the mitochondrial fate of pyruvate, whereas LDH directs its anaerobic fate (Figure 4A). When directed with the ubiquitous *da-gal4* driver, *PK-RNAi* provoked early larval lethality, *PFK1-RNAi* and *PDH-RNAi* provoked larval lethality at L2 or L3 stages, whereas *LDH-RNAi* induced a semi-lethal phenotype at larval or pupal stages (Figure 4B).

Second, we monitored the phosphorylation of the *Drosophila* S6Kinase (dS6K) and Akt as read-out of the activity of mTORC1- and Ilp-signaling respectively. To circumvent the early lethality, the *da-gal4* driver was combined with a ubiquitous thermo-sensitive form of the Gal4 inhibitor, Gal80^ts^ (*tub-gal80*^*ts*^) that blocks Gal4 activity at 21°C but not at 29°C, thereby allowing RNAi expression after temperature shift. Each RNAi was ubiquitously induced at early L1 stage and protein extracts were prepared two days later using late L2 larvae. At this stage the larvae were still viable, although those expressing PK-RNAi did not undergo L2/L3 transition and eventually died (Figure 4B). Western-blotting using these L2 protein extracts revealed that RNAi-knockdown of PFK1, LDH or PDH did not affect Akt or dS6K phosphorylation (Figure 4C). In contrast, PK knockdown strongly decreased dS6K phosphorylation and to a lower extent Akt phosphorylation (Figure 4C). These results indicate that mTORC1 signaling may be affected when knocking down PK, but not when knocking down any other enzyme directly linked to glycolysis. Nonetheless, the lethal phenotype of PK-RNAi larvae occurring at the late L2 stage (Figure 4B) might weaken the larvae, inducing a subsequent effect on mTOR signaling.

To evaluate the requirement of glycolysis for adult survival, RNAi-knockdown was induced by temperature shift to 29°C in newly emerged flies and lethality was counted every second day. In both males and females, PK and PFK1 knockdown provoked lethality between 10 to 14 days after temperature shift (Figure 4D). Knockdown of PDH and LDH also induced adult lethality, although not as soon as PK and PFK1 knockdown (Figure 4D). As a comparison, to evaluate the consequence of disrupting fatty acid synthesis, we knocked-down FASN (Fatty Acid Synthase, Figure 4A) in adults; about a quarter of *FASN-RNAi* flies died between 10 to 14 days, while the others survived nearly as well as control flies (Figure 4D). Taken together, these data indicate that glycolysis is essential for both larval development and adult survival. However, prior to the appearance of the deleterious phenotype, glycolysis knockdown is unlikely to impinge on mTOR signaling.

### Cell-autonomous requirement of glycolysis for Ilp-but not mTORC1-dependent overgrowth

To investigate the requirement of glycolysis to sustain cell-autonomous overgrowth dependent on Ilp- and mTORC1-signaling, *PFK1-RNAi, PK-RNAi, PDH-RNAi* and *LDH-RNAi* were induced in *PTEN*^−*/*−^ or *Rheb*^+^ clones. Except a moderate effect of *PK-RNAi*, clones expressing interfering RNA against these metabolic enzymes did not significantly affect the growth of FB cells (Figure 5A-D and 5M). In combined clones, none of the RNAi affected the growth of *Rheb*^+^ clones (Figure 5E-H and 5M). In contrast, the size of *PTEN*^−*/*−^ clones was significantly decreased when co-expressing RNAi against any of these metabolic enzymes (Figure 5I-M). These findings indicate that both aerobic and anaerobic glycolysis are required to sustain cell-autonomous overgrowth dependent on Ilp signaling. In contrast, reducing glycolysis does not counteract cell-autonomous overgrowth dependent on mTORC1 signaling, suggesting the existence of compensatory pathways.

### Linking Lipogenesis to mTORC1- and Ilp-signaling

Since glycolysis and FA synthesis are tightly connected metabolic pathways (Garrido *et al*. 2015), we investigated whether lipogenesis affects Ilp or mTORC1 signaling. FA synthesis is catalyzed by FASN (Figure 4A). The *Drosophila* genome encodes three *FASN* genes, *FASN1* is ubiquitously expressed but not *FASN2* or *FASN3* (Parvy *et al*. 2012; Chung *et al*. 2014; Wicker-Thomas *et al*. 2015). The deletion of the *FASN1* and *FASN2* tandem (*FASN*^*Δ24-23*^ deletion, hereafter called *FASN*^*1-2*^) results in a lethal phenotype that can be rescued by feeding larvae a lipid-complemented diet (beySD) (Garrido *et al*. 2015; Wicker-Thomas *et al*. 2015). We observed that beySD-rescued *FASN*^*1-2*^ mutant larvae exhibited a delay in development, as measured by the duration of larval development to metamorphosis entry (Figure 6A). Further, when beySD-rescued *FASN*^*1-2*^ mutant larvae were transferred at the L2/L3 larval transition onto a 10% sucrose-supplemented-beySD, only a few of them completed the third larval stage and, after an extreme developmental delay, entered metamorphosis (Figure 6A). Delay in development can be due to a default in ecdysone production that results in giant pupae (Parvy *et al*. 2014) or to impaired mTOR signaling that results in reduced body growth (Montagne *et al*. 1999; Oldham *et al*. 2000). Measurements of prepupal weight revealed that *FASN*^*1-2*^ mutant prepupae exhibited a severe reduction in body weight, whether or not they were supplemented with sucrose (Figure 6B), suggesting a default in mTOR signaling. Therefore, we analyzed the phosphorylation of the *Drosophila* S6Kinase (dS6K) and Akt in protein extracts of late feeding L3 larvae. Western-blotting revealed that the dS6K protein resolved in several bands in *FASN*^*1-2*^ extracts, whereas Akt protein was unchanged (Figure 6C). These results suggest that dS6K but not Akt might be degraded in the *FASN*^*1-2*^ mutant background. In addition, dS6K phosphorylation decreased in *FASN*^*1-2*^ extracts and became barely detectable when *FASN*^*1-2*^ larvae were fed a sucrose-supplemented-beySD (Figure 6C). Conversely, the phosphorylation of Akt was unaffected in larvae fed a beySD, although it was slightly decreased in larvae fed a sucrose-supplemented-beySD (Figure 6C). This finding contrasts with our previous observation showing that FB explants of *FASN*^*1-2*^ mutant larvae were hypersensitive to insulin (Garrido *et al*. 2015). However, *FASN*^*1-2*^ mutants also exhibited a decrease in food intake (Garrido *et al*. 2015), which might induce a systemic suppression of dS6K phosphorylation, while FB explant were cultured in nutrient media supplemented with insulin. Therefore, to determine whether *FASN* mutation affects mTOR signaling at the cell-autonomous level, we analyzed P-S6 and P-Akt in *FASN*^*1-2*^ mutant clones in the FB. As for control clones, about half of the *FASN*^*1-2*^ clonal cells were P-S6 positive (Figure 2J and 2K). Furthermore, no effect on P-Akt was observed in *FASN*^*1-2*^ clonal cells (Figure 2D). In summary, these findings reveal that disrupting FA synthesis does not significantly affect mTORC1 and Ilp signaling at the cell-autonomous level, although it seems to impinge on mTORC1 signaling when inhibited in the whole animal whether directly or indirectly.

### Cell-autonomous requirement of FA synthesis for Ilp-but not mTORC1-dependent overgrowth

To determine, whether lipogenesis is required at the cell-autonomous level to sustain mTORC1 and/or Ilp dependent growth, we analyzed *FASN*^*1-2*^ clones while enhancing either of the mTOR signaling branch in FB cells. We previously reported (Garrido *et al*. 2015) that *FASN*^*1-2*^ clonal cells in the FB were slightly reduced in size and that this effect was dramatically increased in larvae fed a 20%-SSD (Figure S1 and Figure 7M). Therefore, we generated *PTEN*^−*/*−^ and *Rheb*^+^ clones combined or not with the *FASN*^*1-2*^ mutation and analyzed them in the FB of larvae fed either a standard diet or a 20%-SSD. As compared to the standard diet, feeding larvae a 20%-SSD had no effect on the size of *Rheb*^+^ clonal cells, but significantly reduced the size of *PTEN*^−*/*−^ and of *PTEN*^−*/*−^;*Rheb*^+^ clonal cells (Figure 7A-F and 7M). Further, when combined with the *FASN*^*1-2*^ mutation, *PTEN*^−*/*−^ but not *Rheb*^+^ clones were significantly reduced in size (Figure 7G-H and 7M). The *FASN*^*1-2*^ mutation also provoked a severe size reduction of *PTEN*^−*/*−^,*Rheb*^+^ clones (Figure 7I and 7M). Moreover, as compared to the standard diet, feeding larvae a 20%-SSD induced a significant size reduction of *FASN*^*1-2*^;*Rheb*^+^, *FASN*^*1-2*^,*PTEN*^−*/*−^ and *FASN*^*1-2*^,*PTEN*^−*/*−^;*Rheb*^+^ clonal cells (Figure 7J-L and 7M). Of note, except for the *FASN*^*1-2*^;*Rheb*^+^ clonal cells in larvae fed a 20%-SSD that exhibited a size roughly identical to that of the surrounding control cells (Figure 7J), the cell size was always bigger than the controls (Figure 7M). These findings indicate that, in larvae fed a standard diet, FA synthesis is at least in part required to sustain over-growth induced by Ilp, but not mTORC1. They also reveal that additional dietary sucrose is rather detrimental for the growth of cells either deficient for FA synthesis or over-active for Ilp signaling, suggesting that these cells have a restricted homeostatic ability to adjust to an unbalanced diet, whereas mTORC1 activated cells at least in part maintain this ability.

## DISCUSSION

In this study, we used the powerful *Drosophila* genetics to investigate the functional links between the glycolytic/lipogenic axis and mTORC1- or Ilp-dependent growth. In agreement with previous studies (Radimerski *et al*. 2002a; Radimerski *et al*. 2002b; Dong and Pan 2004; Montagne *et al*. 2010; Pallares-Cartes *et al*. 2012), we show that mTORC1 and Ilp signaling work independently in the *Drosophila* FB. Further, we provide evidence that the previously described *mTOR*^*2L1*^ mutation that likely results in a kinase-inactive protein (Oldham *et al*. 2000) affects mTORC1 but not Ilp signaling. Congruently, a study on a *Drosophila rictor* mutant reported that the mTORC2 complex was not required to sustain Akt-dependent growth, but rather to play as a rheostat for this signaling branch (Hietakangas and Cohen 2007). Although this study suggests that mTOR is dispensable for Akt activity, we show that Akt activity and Ilp-dependent overgrowth are suppressed in *mTOR*^*ΔP*^ mutant indicating that the mTOR protein is required for these processes.

On one hand, to mimic the effect that might be induced by drug treatment with a systemic inhibitor, we dampened the glycolytic/lipogenic axis or enhanced mTORC1 or Ilp signaling in the entire organism. On the other hand, to monitor the cell growth process that spans the entire developmental program at the cell-autonomous level, we analyzed clonal FB cells in mosaic animals. Intriguingly, our study reveals apparent contradictory effects between perturbations at the whole body and cell-autonomous levels. At the organismal level, knockdown of glycolytic enzymes or deficiency of FASN result in animal lethality. However, *FASN*^*1-2*^ mutant animals supplemented with dietary lipids can survive but exhibit a dramatic overall growth suppression. This growth defect might result from a decrease in mTORC1 activity that is strongly reduced in *FASN*^*1-2*^ mutant animals, suggesting that mTORC1 but not Ilp signaling relies on lipogenesis. In contrast, at the cell autonomous level, the mutation of *FASN*^*1-2*^ restrains Ilp but not mTORC1 dependent overgrowth in FB cells. These apparent contradictory findings, suggest that the growth defect and the reduction of mTORC1 activity in *FASN*^*1-2*^ mutants are not due to the addition of cell-autonomous effects but rather to a systemic regulation. Potentially, FASN default might affect the activity of a specific tissue, as for instance, the neurosecretory cells that synthesize and secrete Ilps, which promote systemic body growth (Rulifson *et al*. 2002). Alternatively, considering that mTORC1 directly responds to nutrients (Dibble and Manning 2013; Groenewoud and Zwartkruis 2013; Montagne 2016), the drop of mTORC1 activity may be a consequence of feeding, since we previously reported a decrease in nutrient uptake in *FASN*^*1-2*^ mutant animals (Garrido *et al*. 2015). Consistently, a previous study on the transcription factor Mondo — the *Drosophila* homologue of mondoA and ChREBP that regulate the glycolytic/lipogenic axis in response to dietary sugar (Mattila *et al*. 2015; Richards *et al*. 2017)— suggests the existence of a FASN-dependent effect in the FB on the control of food intake (Sassu *et al*. 2012). FB-knockdown of *mondo* results in the lack of sucrose-induced expression of *FASN1* and in a decrease in food intake. This study suggests that the FASN default perturbs body homeostasis and indirectly affects the neuronal control of feeding behavior. However, it does not exclude that a lipogenic defect in neuronal cells may also directly impinge on feeding behavior. Finally, the drop of mTORC1 activity observed in *FASN*^*1-2*^ mutants may be a consequence of malonyl-CoA accumulation, since mTOR malonylation has been reported to inhibit mTORC1- but not Ilp/mTORC2-dependent activity (Bruning *et al*. 2018). Malonylation of mTOR may also account for the size reduction of *FASN*^*1-2*^ mutant cells over-expressing Rheb in animals fed a 20%-SSD, consistent with the increased expression of lipogenic enzymes induced by dietary sucrose (Garrido *et al*. 2015). Thus, mTOR malonylation and the subsequent decrease in mTORC1 activity might occur only when interfering with a context of high demand for lipogenesis, an issue that should be investigated in the future.

Our study reveals that over-activation of mTORC1 and to a lesser extent of Ilp signaling, results in a decrease in glycogen and TAG stores and in circulating trehalose, suggesting that activation of either signaling branch enhances metabolite consumption to sustain cell growth. It is therefore surprising that activation of neither mTORC1 nor Ilp signaling induces an increase in body weight. Nonetheless, overall body growth depends on an intricate regulatory network that integrates cell-autonomous effects and humoral messages. For instance, previous studies reported that activation of Ilp signaling within the ring gland, results in a systemic decrease in body growth (Caldwell *et al*. 2005; Colombani *et al*. 2005; Mirth 2005). Therefore, ubiquitous activation of mTORC1 or Ilp signaling is likely to promote the growth of most cells but might concurrently perturb endocrine signals dampening overall growth. Of note, we observed that larvae fed a 20%-SSD result in pupae with reduced body weight, an effect that is partially suppressed when either mTORC1 or Ilp signaling is over-activated. The fact that the overall body weight of these animals is maintained within a range likely compatible with organismal survival contrasts with the observed high rate of lethality. The decrease in stores and circulating sugars suggests that in these animals each cell tends to increase its basal metabolism evoking an egoist behavior that might perturb the equilibrium between cell-autonomous and systemic regulation. Thus, in a stressful situation, as when animals are fed a 20%-SSD, the need of a tight adjustment to an unbalanced diet may enhance the distortion between cell-autonomous effects and systemic regulation, resulting in an increased rate of lethality.

A plethora of studies in mammalian cells indicate that mTOR activation directs metabolism towards glucose consumption, storage and anabolism (Gottlob *et al*. 2001; Inoki *et al*. 2003b; Hahn-Windgassen *et al*. 2005; Duvel *et al*. 2010; Peterson *et al*. 2011; Houddane *et al*. 2017; Jaldin-Fincati *et al*. 2017; Wipperman *et al*. 2019). Our study rather suggests that in the *Drosophila* larvae, mTOR promotes metabolite consumption through glycolysis but not storage. However, at the cell-autonomous level, we observe that inhibition of lipogenesis or glycolysis restrains neither larval FB cell growth nor overgrowth induced by mTORC1 stimulation in these cells. These findings counteract the idea that mTORC1 potentiates a glycolytic/lipogenic axis (Duvel *et al*. 2010) to sustain cell growth. To overcome the lack of glycolytic products and of membrane lipids, these cells may benefit of a transfer from neighboring cells and might favor alternative metabolic pathways, including glutamine catabolism to feed TCA anaplerosis, which has been shown to be a crucial pathway in mTORC1-stimulated mammalian cells (Choo *et al*. 2010; Csibi *et al*. 2013; Csibi *et al*. 2014). Nonetheless, such compensatory processes do not fully operate to sustain Ilp-dependent overgrowth. In these cells, the mutation of PTEN potentially impedes the ability to modulate this signaling branch. Therefore, it is tempting to speculate that the modulation of Ilp signaling at least in part contributes to the regulation of these compensatory processes.

As a coordinator of growth and metabolism, mTOR plays a central role in tumor development (Dowling *et al*. 2010; Harachi *et al*. 2018; Mossmann *et al*. 2018; Tian *et al*. 2019). PTEN, the tumor suppressor that counteracts PI3K activity downstream of the Ilp receptor, is deficient in several human cancers (Cully *et al*. 2006). Mutation of TSC1 or TSC2, which results in mTORC1 hyper-activation, is associated with benign tumors but also with brain, kidney and lung destructive diseases (Henske *et al*. 2016). To investigate the role of mTOR regarding tumor development, a recent study reported the generation of liver-specific double knockout mice for TSC1 and PTEN (Guri *et al*. 2017). These mice develop hepatic steatosis that eventually progresses to hepatocellular carcinoma. Both processes are suppressed in mice fed the mTORC1/2 inhibitors INK128, but not the mTORC1 inhibitor rapamycin, supporting an Ilp/mTORC2 specific effect. The combination of inhibitors against mTOR and metabolism is currently under clinical investigation to fight cancers (Mossmann *et al*. 2018). Importantly, our study reveals that ubiquitous inhibition of basal metabolism produces dramatic effects during development, while at the cell-autonomous level, it only moderates growth induced by over-activation of Ilp/mTORC2 signaling. Therefore, the use of drug therapy to fight cancer must be taken with caution, in particular if organismal development is not complete and most efforts should be made to selectively target sick tissues.

## AKNOWLEDGMENTS

We wish to thank D Petit for preparing the fly media, H Stocker for fly stocks, A Teleman for the phospho-S6 antibody, M Gettings for editing the manuscript, and the NIG and VDRC stock centers for RNAi fly strains. We wish to thank the *French government* for fellowship to MD (MENRT 2015-155) and DG (MRT 2011-78), the *Fondation pour la Recherche Médicale* for fellowship to DG (FDT201 4093 0800), the *Fondation ARC* for grant support to JM (projet ARC 1555286) and the *Ligue de Recherche contre le Cancer* for grant support to JM (M27218). The authors declare no competing financial interests.

## AUTHOR CONTRIBUTIONS

JM designed the experiments; MD, DG, MP, TR and JM performed the experiments; MD, DG, ALR and JM analyzed the results; and JM wrote the manuscript.

## LITERATURE CITED

Antikainen, H., M. Driscoll, G. Haspel and R. Dobrowolski, 2017 TOR-mediated regulation of metabolism in aging. Aging Cell 16: 1219–1233.

Bates, D., M. Mächler, B. M. Bolker and S. Walker, 2015 Fitting Linear Mixed-Effects Models Using lme4. Journal of Statistical Software 67: 1–48.

Bohni, R., J. Riesgo-Escovar, S. Oldham, W. Brogiolo, H. Stocker et al., 1999 Autonomous control of cell and organ size by CHICO, a Drosophila homolog of vertebrate IRS1-4. Cell 97: 865–875.

Bruning, U., F. Morales-Rodriguez, J. Kalucka, J. Goveia, F. Taverna et al., 2018 Impairment of Angiogenesis by Fatty Acid Synthase Inhibition Involves mTOR Malonylation. Cell Metab 28: 866–880 e815.

Caldwell, P. E., M. Walkiewicz and M. Stern, 2005 Ras activity in the Drosophila prothoracic gland regulates body size and developmental rate via ecdysone release. Curr Biol 15: 1785–1795.

Caron, A., D. Richard and M. Laplante, 2015 The Roles of mTOR Complexes in Lipid Metabolism. Annu Rev Nutr 35: 321–348.

Choo, A. Y., S. G. Kim, M. G. Vander Heiden, S. J. Mahoney, H. Vu et al., 2010 Glucose addiction of TSC null cells is caused by failed mTORC1-dependent balancing of metabolic demand with supply. Mol Cell 38: 487–499.

Chung, H., D. W. Loehlin, H. D. Dufour, K. Vaccarro, J. G. Millar et al., 2014 A single gene affects both ecological divergence and mate choice in Drosophila. Science 343: 1148–1151.

Colombani, J., L. Bianchini, S. Layalle, E. Pondeville, C. Dauphin-Villemant et al., 2005 Antagonistic actions of ecdysone and insulins determine final size in Drosophila. Science 310: 667–670.

Csibi, A., S. M. Fendt, C. Li, G. Poulogiannis, A. Y. Choo et al., 2013 The mTORC1 pathway stimulates glutamine metabolism and cell proliferation by repressing SIRT4. Cell 153: 840–854.

Csibi, A., G. Lee, S. O. Yoon, H. Tong, D. Ilter et al., 2014 The mTORC1/S6K1 pathway regulates glutamine metabolism through the eIF4B-dependent control of c-Myc translation. Curr Biol 24: 2274–2280.

Cully, M., H. You, A. J. Levine and T. W. Mak, 2006 Beyond PTEN mutations: the PI3K pathway as an integrator of multiple inputs during tumorigenesis. Nat Rev Cancer 6: 184–192.

Dibble, C. C., W. Elis, S. Menon, W. Qin, J. Klekota et al., 2012 TBC1D7 is a third subunit of the TSC1-TSC2 complex upstream of mTORC1. Mol Cell 47: 535–546.

Dibble, C. C., and B. D. Manning, 2013 Signal integration by mTORC1 coordinates nutrient input with biosynthetic output. Nat Cell Biol 15: 555–564.

Dietzl, G., D. Chen, F. Schnorrer, K. C. Su, Y. Barinova et al., 2007 A genome-wide transgenic RNAi library for conditional gene inactivation in Drosophila. Nature 448: 151–156.

Dong, J., and D. Pan, 2004 Tsc2 is not a critical target of Akt during normal Drosophila development. Genes Dev 18: 2479–2484.

Dowling, R. J., I. Topisirovic, B. D. Fonseca and N. Sonenberg, 2010 Dissecting the role of mTOR: lessons from mTOR inhibitors. Biochim Biophys Acta 1804: 433–439.

Duvel, K., J. L. Yecies, S. Menon, P. Raman, A. I. Lipovsky et al., 2010 Activation of a metabolic gene regulatory network downstream of mTOR complex 1. Mol Cell 39: 171–183.

Edgar, B. A., and T. L. Orr-Weaver, 2001 Endoreplication cell cycles: more for less. Cell 105: 297–306.

Engelman, J. A., J. Luo and L. C. Cantley, 2006 The evolution of phosphatidylinositol 3-kinases as regulators of growth and metabolism. Nat Rev Genet 7: 606–619.

Garami, A., F. J. Zwartkruis, T. Nobukuni, M. Joaquin, M. Roccio et al., 2003 Insulin activation of Rheb, a mediator of mTOR/S6K/4E-BP signaling, is inhibited by TSC1 and 2. Mol Cell 11: 1457–1466.

Garrido, D., T. Rubin, M. Poidevin, B. Maroni, A. Le Rouzic et al., 2015 Fatty Acid Synthase Cooperates with Glyoxalase 1 to Protect against Sugar Toxicity. PLoS Genet 11: e1004995.

Goberdhan, D. C., M. H. Ogmundsdottir, S. Kazi, B. Reynolds, S. M. Visvalingam et al., 2009 Amino acid sensing and mTOR regulation: inside or out? Biochem Soc Trans 37: 248–252.

Gottlob, K., N. Majewski, S. Kennedy, E. Kandel, R. B. Robey et al., 2001 Inhibition of early apoptotic events by Akt/PKB is dependent on the first committed step of glycolysis and mitochondrial hexokinase. Genes Dev 15: 1406–1418.

Groenewoud, M. J., and F. J. Zwartkruis, 2013 Rheb and mammalian target of rapamycin in mitochondrial homoeostasis. Open Biol 3: 130185.

Guri, Y., M. Colombi, E. Dazert, S. K. Hindupur, J. Roszik et al., 2017 mTORC2 Promotes Tumorigenesis via Lipid Synthesis. Cancer Cell 32: 807–823 e812.

Haeusler, R. A., T. E. McGraw and D. Accili, 2018 Biochemical and cellular properties of insulin receptor signalling. Nat Rev Mol Cell Biol 19: 31–44.

Hagiwara, A., M. Cornu, N. Cybulski, P. Polak, C. Betz et al., 2012 Hepatic mTORC2 activates glycolysis and lipogenesis through Akt, glucokinase, and SREBP1c. Cell Metab 15: 725–738.

Hahn-Windgassen, A., V. Nogueira, C. C. Chen, J. E. Skeen, N. Sonenberg et al., 2005 Akt activates the mammalian target of rapamycin by regulating cellular ATP level and AMPK activity. J Biol Chem 280: 32081–32089.

Harachi, M., K. Masui, Y. Okamura, R. Tsukui, P. S. Mischel et al., 2018 mTOR Complexes as a Nutrient Sensor for Driving Cancer Progression. Int J Mol Sci 19.

Hay, N., and N. Sonenberg, 2004 Upstream and downstream of mTOR. Genes Dev 18: 1926–1945.

Henske, E. P., S. Jozwiak, J. C. Kingswood, J. R. Sampson and E. A. Thiele, 2016 Tuberous sclerosis complex. Nat Rev Dis Primers 2: 16035.

Hietakangas, V., and S. M. Cohen, 2007 Re-evaluating AKT regulation: role of TOR complex 2 in tissue growth. Genes Dev 21: 632–637.

Holm, S., 1079 A simple sequentially rejective multiple test procedure. Scand J Statist 6.

Hothorn, T., F. Bretz and P. Westfall, 2008 Simultaneous inference in general parametric models. Biom J 50: 346–363.

Houddane, A., L. Bultot, L. Novellasdemunt, M. Johanns, M. A. Gueuning et al., 2017 Role of Akt/PKB and PFKFB isoenzymes in the control of glycolysis, cell proliferation and protein synthesis in mitogen-stimulated thymocytes. Cell Signal 34: 23–37.

Howell, J. J., S. J. Ricoult, I. Ben-Sahra and B. D. Manning, 2013 A growing role for mTOR in promoting anabolic metabolism. Biochem Soc Trans 41: 906–912.

Inoki, K., Y. Li, T. Xu and K. L. Guan, 2003a Rheb GTPase is a direct target of TSC2 GAP activity and regulates mTOR signaling. Genes Dev 17: 1829–1834.

Inoki, K., T. Zhu and K. L. Guan, 2003b TSC2 mediates cellular energy response to control cell growth and survival. Cell 115: 577–590.

Jaldin-Fincati, J. R., M. Pavarotti, S. Frendo-Cumbo, P. J. Bilan and A. Klip, 2017 Update on GLUT4 Vesicle Traffic: A Cornerstone of Insulin Action. Trends Endocrinol Metab 28: 597–611.

Kim, D. H., D. D. Sarbassov, S. M. Ali, J. E. King, R. R. Latek et al., 2002 mTOR interacts with raptor to form a nutrient-sensitive complex that signals to the cell growth machinery. Cell 110: 163–175.

Lamming, D. W., and D. M. Sabatini, 2013 A Central role for mTOR in lipid homeostasis. Cell Metab 18: 465–469.

Laplante, M., and D. M. Sabatini, 2012 mTOR signaling in growth control and disease. Cell 149: 274–293.

Lee, T., and L. Luo, 2001 Mosaic analysis with a repressible cell marker (MARCM) for Drosophila neural development. Trends Neurosci 24: 251–254.

Lehmann, M., 2018 Endocrine and physiological regulation of neutral fat storage in Drosophila. Mol Cell Endocrinol 461: 165–177.

Lien, E. C., C. C. Dibble and A. Toker, 2017 PI3K signaling in cancer: beyond AKT. Curr Opin Cell Biol 45: 62–71.

Ma, X. M., and J. Blenis, 2009 Molecular mechanisms of mTOR-mediated translational control. Nat Rev Mol Cell Biol 10: 307–318.

Magnuson, B., B. Ekim and D. C. Fingar, 2012 Regulation and function of ribosomal protein S6 kinase (S6K) within mTOR signalling networks. Biochem J 441: 1–21.

Majewski, N., V. Nogueira, P. Bhaskar, P. E. Coy, J. E. Skeen et al., 2004 Hexokinase-mitochondria interaction mediated by Akt is required to inhibit apoptosis in the presence or absence of Bax and Bak. Mol Cell 16: 819–830.

Mattila, J., E. Havula, E. Suominen, M. Teesalu, I. Surakka et al., 2015 Mondo-Mlx Mediates Organismal Sugar Sensing through the Gli-Similar Transcription Factor Sugarbabe. Cell Rep 13: 350–364.

McManus, E. J., K. Sakamoto, L. J. Armit, L. Ronaldson, N. Shpiro et al., 2005 Role that phosphorylation of GSK3 plays in insulin and Wnt signalling defined by knockin analysis. EMBO J 24: 1571–1583.

Mirth, C., 2005 Ecdysteroid control of metamorphosis in the differentiating adult leg structures of Drosophila melanogaster. Dev Biol 278: 163–174.

Montagne, J., 2016 A Wacky Bridge to mTORC1 Dimerization. Dev Cell 36: 129–130.

Montagne, J., C. Lecerf, J. P. Parvy, J. M. Bennion, T. Radimerski et al., 2010 The nuclear receptor DHR3 modulates dS6 kinase-dependent growth in Drosophila. PLoS Genet 6: e1000937.

Montagne, J., T. Radimerski and G. Thomas, 2001 Insulin signaling: lessons from the Drosophila tuberous sclerosis complex, a tumor suppressor. Sci STKE 2001: PE36.

Montagne, J., M. J. Stewart, H. Stocker, E. Hafen, S. C. Kozma et al., 1999 Drosophila S6 kinase: a regulator of cell size. Science 285: 2126–2129.

Montagne, J., and G. Thomas, 2004 S6K integrates nutrient and mitogen signals to control cell growth., pp. 265–298 in Cell growth: control of cell size., edited by M. Hall, Raff, M., Thomas, G. Cold Spring Harbor Press.

Morata, G., and P. Ripoll, 1975 Minutes: mutants of drosophila autonomously affecting cell division rate. Dev Biol 42: 211–221.

Mossmann, D., S. Park and M. N. Hall, 2018 mTOR signalling and cellular metabolism are mutual determinants in cancer. Nat Rev Cancer 18: 744–757.

Nakae, J., T. Kitamura, D. L. Silver and D. Accili, 2001 The forkhead transcription factor Foxo1 (Fkhr) confers insulin sensitivity onto glucose-6-phosphatase expression. J Clin Invest 108: 1359–1367.

Oldham, S., J. Montagne, T. Radimerski, G. Thomas and E. Hafen, 2000 Genetic and biochemical characterization of dTOR, the Drosophila homolog of the target of rapamycin. Genes Dev 14: 2689–2694.

Padmanabha, D., and K. D. Baker, 2014 Drosophila gains traction as a repurposed tool to investigate metabolism. Trends Endocrinol Metab 25: 518–527.

Pallares-Cartes, C., G. Cakan-Akdogan and A. A. Teleman, 2012 Tissue-specific coupling between insulin/IGF and TORC1 signaling via PRAS40 in Drosophila. Dev Cell 22: 172–182.

Parvy, J. P., L. Napal, T. Rubin, M. Poidevin, L. Perrin et al., 2012 Drosophila melanogaster Acetyl-CoA-carboxylase sustains a fatty acid-dependent remote signal to waterproof the respiratory system. PLoS Genet 8: e1002925.

Parvy, J. P., P. Wang, D. Garrido, A. Maria, C. Blais et al., 2014 Forward and feedback regulation of cyclic steroid production in Drosophila melanogaster. Development 141: 3955–3965.

Peterson, T. R., S. S. Sengupta, T. E. Harris, A. E. Carmack, S. A. Kang et al., 2011 mTOR complex 1 regulates lipin 1 localization to control the SREBP pathway. Cell 146: 408–420.

Polak, P., N. Cybulski, J. N. Feige, J. Auwerx, M. A. Ruegg et al., 2008 Adipose-specific knockout of raptor results in lean mice with enhanced mitochondrial respiration. Cell Metab 8: 399–410.

Radimerski, T., J. Montagne, M. Hemmings-Mieszczak and G. Thomas, 2002a Lethality of Drosophila lacking TSC tumor suppressor function rescued by reducing dS6K signaling. Genes Dev 16: 2627–2632.

Radimerski, T., J. Montagne, F. Rintelen, H. Stocker, J. van der Kaay et al., 2002b dS6K-regulated cell growth is dPKB/dPI(3)K-independent, but requires dPDK1. Nat Cell Biol 4: 251–255.

Richards, P., S. Ourabah, J. Montagne, A. F. Burnol, C. Postic et al., 2017 MondoA/ChREBP: The usual suspects of transcriptional glucose sensing; Implication in pathophysiology. Metabolism 70: 133–151.

Robey, R. B., and N. Hay, 2009 Is Akt the “Warburg kinase”-Akt-energy metabolism interactions and oncogenesis. Semin Cancer Biol 19: 25–31.

Romero-Pozuelo, J., C. Demetriades, P. Schroeder and A. A. Teleman, 2017 CycD/Cdk4 and Discontinuities in Dpp Signaling Activate TORC1 in the Drosophila Wing Disc. Dev Cell 42: 376–387 e375.

Rulifson, E. J., S. K. Kim and R. Nusse, 2002 Ablation of insulin-producing neurons in flies: growth and diabetic phenotypes. Science 296: 1118–1120.

Sarbassov, D. D., D. A. Guertin, S. M. Ali and D. M. Sabatini, 2005 Phosphorylation and regulation of Akt/PKB by the rictor-mTOR complex. Science 307: 1098–1101.

Sassu, E. D., J. E. McDermott, B. J. Keys, M. Esmaeili, A. C. Keene et al., 2012 Mio/dChREBP coordinately increases fat mass by regulating lipid synthesis and feeding behavior in Drosophila. Biochem Biophys Res Commun 426: 43–48.

Saxton, R. A., and D. M. Sabatini, 2017 mTOR Signaling in Growth, Metabolism, and Disease. Cell 168: 960–976.

Shimobayashi, M., and M. N. Hall, 2014 Making new contacts: the mTOR network in metabolism and signalling crosstalk. Nat Rev Mol Cell Biol 15: 155–162.

Stocker, H., T. Radimerski, B. Schindelholz, F. Wittwer, P. Belawat et al., 2003 Rheb is an essential regulator of S6K in controlling cell growth in Drosophila. Nat Cell Biol 5: 559–565.

Tian, T., X. Li and J. Zhang, 2019 mTOR Signaling in Cancer and mTOR Inhibitors in Solid Tumor Targeting Therapy. Int J Mol Sci 20.

Ugur, B., K. Chen and H. J. Bellen, 2016 Drosophila tools and assays for the study of human diseases. Dis Model Mech 9: 235–244.

Wangler, M. F., Y. Hu and J. M. Shulman, 2017 Drosophila and genome-wide association studies: a review and resource for the functional dissection of human complex traits. Dis Model Mech 10: 77–88.

Wicker-Thomas, C., D. Garrido, G. Bontonou, L. Napal, N. Mazuras et al., 2015 Flexible origin of hydrocarbon/pheromone precursors in Drosophila melanogaster. J Lipid Res 56: 2094–2101.

Wipperman, M. F., D. C. Montrose, A. M. Gotto, Jr. and D. P. Hajjar, 2019 Mammalian Target of Rapamycin: A Metabolic Rheostat for Regulating Adipose Tissue Function and Cardiovascular Health. Am J Pathol 189: 492–501.

Yang, H., X. Jiang, B. Li, H. J. Yang, M. Miller et al., 2017 Mechanisms of mTORC1 activation by RHEB and inhibition by PRAS40. Nature 552: 368–373.

Yuan, M., E. Pino, L. Wu, M. Kacergis and A. A. Soukas, 2012 Identification of Akt-independent regulation of hepatic lipogenesis by mammalian target of rapamycin (mTOR) complex 2. J Biol Chem 287: 29579–29588.

Zhang, H., J. P. Stallock, J. C. Ng, C. Reinhard and T. P. Neufeld, 2000 Regulation of cellular growth by the Drosophila target of rapamycin dTOR. Genes Dev 14: 2712–2724.

