## Supplementary material for "Differential Metabolic Sensitivity of Insulin-like-response- and mTORC1-Dependent Overgrowth in *Drosophila* Fat Cells": Figure S1

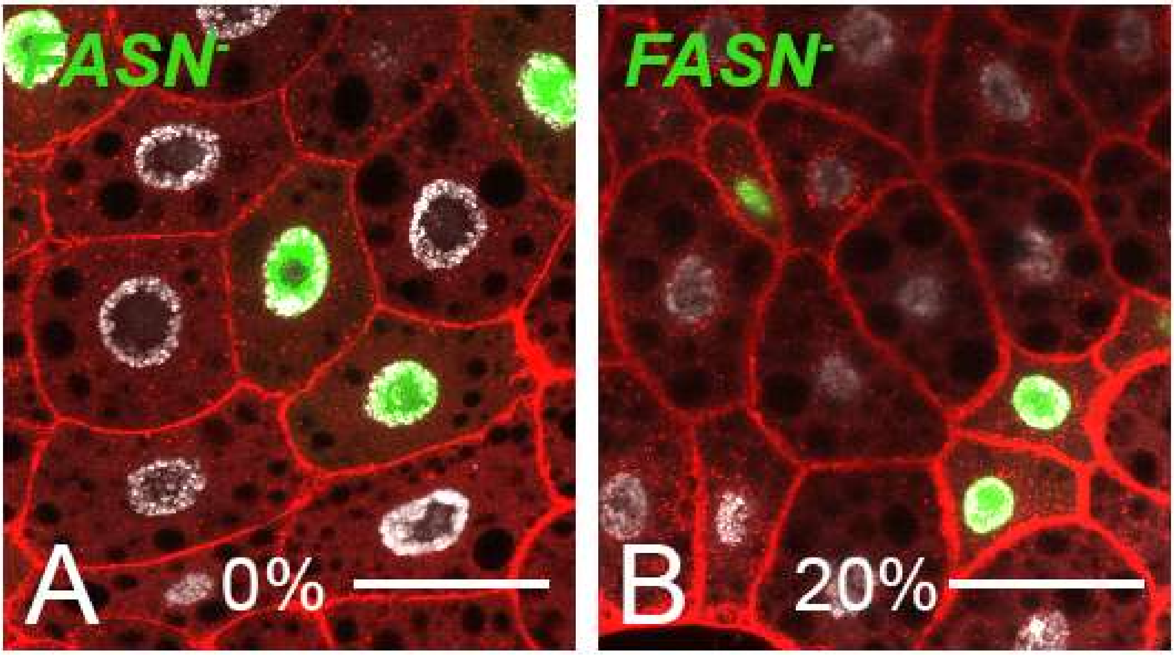


**Figure S1. Effect of dietary sugar on *FASN*$^{1\text{-}2}$ mutant cells.** *FASN*$^{1\text{-}2}$ MARCM clones labeled by GFP (green) in the FB of L3 larvae fed either a standard (A) or a 20%-SSD (B). Nuclei are labeled with DAPI (silver) and membranes by phalloidin (red). Scale bars: 50µm.
