## Supplementary material for "Differential Metabolic Sensitivity of Insulin-like-response- and mTORC1-Dependent Overgrowth in *Drosophila* Fat Cells": Figure S2

**A**

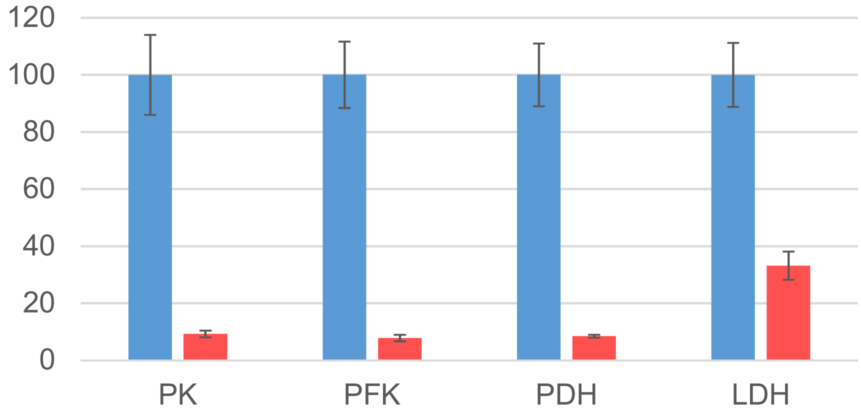


**B**

| | | |
|---|---|---|
| PFK1: | 5'-CCAGTGCCTTCGATCGTATT-3' | 5'-GTTGCCGTCCAGAGAGATG-3' |
| PK: | 5'-GAACGTAACCATCTACGATGAGG-3' | 5'- CATGTGCTCCAGTTGGGTAT-3' |
| LDH: | 5'-CGTCTCCACCTCCGTTT-3' | 5'-TCACACCGTTGGCATTGA-3' |
| PDH: | 5'-TGATCCCATCACCTCCTTCA-3' | 5'-CCTTCAGATCGATGGCCTTAAC-3' |

**Figure S2. Knockdown efficacy of the RNAi to the glycolytic enzymes.** (**A**) mRNA expression in control larvae (blue bars, +/- standard error) was adjusted to 100. mRNA levels after RNAi knockdown (red bars, +/- standard error) was calculated as a percentage of corresponding expression in control larvae. (**B**) Oligonucleotides used in RT-Q-PCR for each glycolytic enzyme.
