## Supplementary material for "Differential Metabolic Sensitivity of Insulin-like-response- and mTORC1-Dependent Overgrowth in *Drosophila* Fat Cells": Table S1

| Figure 1M | Estimate | Std. error | Adj. P-value |
|---|---|---|---|
| *Rheb*$^{+}$ vs. Co | 0.4584 | 0.0715 | 0.0000 |
| *PTEN*$^{-}$ vs. Co | 1.0184 | 0.0644 | 0.0000 |
| *PTEN*$^{-}$;*Rheb*$^{+}$ vs. Co | 1.4573 | 0.0603 | 0.0000 |
| *PTEN*$^{-}$;*Rheb*$^{+}$ vs. *Rheb*$^{+}$ | 0.9989 | 0.0685 | 0.0000 |
| *PTEN*$^{-}$;*Rheb*$^{+}$ vs. *PTEN*$^{-}$ | 0.4389 | 0.0611 | 0.0000 |

| Figure 5M | Estimate | Std. error | Adj. P-value |
|---|---|---|---|
| *PFK1i* vs. Co | -0.0533 | 0.0672 | 1.0000 |
| *PKi* vs. Co | -0.1932 | 0.0677 | 0.0343 |
| *LDHi* vs. Co | -0.0665 | 0.0671 | 1.0000 |
| *PDHi* vs. Co | -0.0679 | 0.0706 | 1.0000 |
| *PTEN*$^{-}$;*PFK1i* vs. *PTEN*$^{-}$ | -0.2100 | 0.0673 | 0.0162 |
| *PTEN*$^{-}$;*PKi* vs. *PTEN*$^{-}$ | -0.3699 | 0.0677 | 0.0000 |
| *PTEN*$^{-}$;*LDHi* vs. *PTEN*$^{-}$ | -0.3726 | 0.0683 | 0.0000 |
| *PTEN*$^{-}$;*PDHi* vs. *PTEN*$^{-}$ | -0.2566 | 0.0652 | 0.0008 |
| *Rheb*$^{+}$,*PFK1i* vs. *Rheb*$^{+}$ | 0.0384 | 0.0773 | 1.0000 |
| *Rheb*$^{+}$,*PKi* vs. *Rheb*$^{+}$ | 0.1466 | 0.0675 | 0.1796 |
| *Rheb*$^{+}$,*LDHi* vs. *Rheb*$^{+}$ | 0.0612 | 0.0746 | 1.0000 |
| *Rheb*$^{+}$,*PDHi* vs. *Rheb*$^{+}$ | -0.1640 | 0.0702 | 0.1371 |

| Figure 7M | Estimate | Std. error | Adj. P-value |
|---|---|---|---|
| *FASN*$^{-}$ 0% vs. 20% | -0.5209 | 0.0709 | 0.0000 |
| *Rheb*$^{+}$ 0% vs. 20% | 0.0259 | 0.0907 | 1.0000 |
| *FASN*$^{-}$;*Rheb*$^{+}$ 0% vs. 20% | -0.6090 | 0.0645 | 0.0000 |
| *PTEN*$^{-}$ 0% vs. 20% | -0.3326 | 0.0736 | 0.0000 |
| *FASN*$^{-}$,*PTEN*$^{-}$ 0% vs. 20% | -0.2753 | 0.0544 | 0.0000 |
| *PTEN*$^{-}$;*Rheb*$^{+}$ 0% vs. 20% | -0.5722 | 0.0722 | 0.0000 |
| FASN-,*PTEN*$^{-}$;*Rheb*$^{+}$ 0% vs. 20% | -0.3094 | 0.0649 | 0.0000 |
| *FASN*$^{-}$,*PTEN*$^{-}$ vs. *FASN*$^{-}$ | 1.0242 | 0.0692 | 0.0000 |
| *FASN*$^{-}$;*Rheb*$^{+}$ vs. *FASN*$^{-}$ | 0.8141 | 0.0781 | 0.0000 |
| *FASN*$^{-}$;*Rheb*$^{+}$ vs. *Rheb*$^{+}$ | -0.0347 | 0.0772 | 1.0000 |
| *FASN*$^{-}$,*PTEN*$^{-}$ vs. PTEN- | -0.3846 | 0.0607 | 0.0000 |
| *FASN*$^{-}$,*PTEN*$^{-}$;*Rheb*$^{+}$ vs. *PTEN*$^{-}$;*Rheb*$^{+}$ | -0.5447 | 0.0526 | 0.0000 |

**Table S1. Statistical tests corresponding to Figures 1M (top), 5M (middle), and 7M (bottom).** The model tests for the difference is the log ratio of *GFP*$^{+}$*/GFP*$^{-}$*surrounding* cells between pairs of genotypes (Co = control), accounting for larvae and series random effects (see the Methods section). P-values were corrected for multiple testing (Holm-Bonferroni method) and quantify the risk of rejecting the true null hypothesis at the table level.
