## Supplementary material for "Differential Metabolic Sensitivity of Insulin-like-response- and mTORC1-Dependent Overgrowth in *Drosophila* Fat Cells": Table S2

| | Effect | Std. error | Adj. P-value |
|---|---|---|---|
| *FASN*$^{-}$ vs. Co | -0.183 | 0.527 | 0.7287 |
| *PTEN*$^{-}$ vs. Co | 0.717 | 0.489 | 0.2857 |
| *Rheb*$^{+}$ vs. Co | 4.126 | 0.879 | 0.0000 |

**Table S2. Statistical test of the difference in P-S6 frequency (Figure 2K) between MARCM clones to control.** The model is a mixed effect, generalized linear model considering frequencies as binomial data and genotype as a fixed effect, while larvae / series were random effects. P-values were adjusted for multiple testing by a Holm-Bonferroni correction.
