## Supplementary material for "Differential Metabolic Sensitivity of Insulin-like-response- and mTORC1-Dependent Overgrowth in *Drosophila* Fat Cells": Table S3

| **A** (Males weight) | **Effect** | **Std. error** | **Adj. P-value** |
|---|---|---|---|
| 0%: *PTENi* vs. Co | 0.0077 | 0.0257 | 0.7653 |
| 0%: *Rheb*$^{++}$ vs. Co | 0.0584 | 0.0238 | 0.0295 |
| 20%: *PTENi* vs. Co | 0.2392 | 0.0236 | 0.0000 |
| 20%: *Rheb*$^{++}$ vs. Co | 0.2241 | 0.0249 | 0.0000 |
| **B** (Female weight) | **Effect** | **Std. error** | **Adj. P-value** |
| 0%: *PTENi* vs. Co | 0.0591 | 0.0351 | 0.1879 |
| 0%: *Rheb*$^{++}$ vs. Co | 0.0107 | 0.0315 | 0.7347 |
| 20%: *PTENi* vs. Co | 0.2357 | 0.0358 | 0.0000 |
| 20%: *Rheb*$^{++}$ vs. Co | 0.3197 | 0.0320 | 0.0000 |
| **C** (Protein) | **Effect** | **Std. error** | **Adj. P-value** |
| 0%: *PTENi* vs. Co | 0.0007 | 0.0004 | 0.5313 |
| 0%: *Rheb*$^{++}$ vs. Co | -0.0002 | 0.0004 | 1.0000 |
| 20%: *PTENi* vs. Co | -0.0002 | 0.0004 | 1.0000 |
| 20%: *Rheb*$^{++}$ vs. Co | -0.0007 | 0.0004 | 0.5313 |
| **D** (TAG) | **Effect** | **Std. error** | **Adj. P-value** |
| 0%: *PTENi* vs. Co | 0.0006 | 0.0044 | 1.0000 |
| 0%: *Rheb*$^{++}$ vs. Co | -0.0118 | 0.0044 | 0.0417 |
| 20%: *PTENi* vs. Co | 0.0004 | 0.0043 | 1.0000 |
| 20%: *Rheb*$^{++}$ vs. Co | -0.0146 | 0.0043 | 0.0084 |
| **E** (Glycogen) | **Effect** | **Std. error** | **Adj. P-value** |
| 0%: *PTENi* vs. Co | -0.8275 | 0.3703 | 0.0370 |
| 0%: *Rheb*$^{++}$ vs. Co | -1.6765 | 0.3703 | 0.0006 |
| 20%: *PTENi* vs. Co | -1.3188 | 0.3903 | 0.0060 |
| 20%: *Rheb*$^{++}$ vs. Co | -1.8334 | 0.3903 | 0.0006 |
| **F** (Trealose) | **Effect** | **Std. error** | **Adj. P-value** |
| 0%: *PTENi* vs. Co | -2.8836 | 0.4507 | 0.0000 |
| 0%: *Rheb*$^{++}$ vs. Co | -2.3548 | 0.4507 | 0.0000 |
| 20%: *PTENi* vs. Co | -3.3097 | 0.4750 | 0.0000 |
| 20%: *Rheb*$^{++}$ vs. Co | -3.2726 | 0.4750 | 0.0000 |

**Table S3. Statistical tests corresponding to Figure 3.** Differences in male (A) or female (B) weight, in protein (C), in TAG (D), in glycogen (E) and in trehalose (F) between control (Co) and prepupae that ubiquitously express either Rheb (*Rheb*$^{++}$) or an RNAi to PTEN (*PTENi*). The column at the right indicates P-values corrected for multiple testing (Holm-Bonferoni correction).
