## Supplementary material for "Differential Metabolic Sensitivity of Insulin-like-response- and mTORC1-Dependent Overgrowth in *Drosophila* Fat Cells": Table S4

| | Estimate | Std. error | t value | Pr(>\|t\|) |
|---|---|---|---|---|
| males, $w^{-}$ : 0% vs. 10% | 0.109 | 0.032 | 3.381 | 0.00162 |
| females, $w^{-}$ : 0% vs. 10% | 0.115 | 0.032 | 3.647 | 0.00093 |
| males, $FASN^{1\text{-}2}$ : 0% vs. 10% | 0.098 | 0.031 | 3.141 | 0.00183 |
| females, $FASN^{1\text{-}2}$ : 0% vs. 10% | 0.124 | 0.031 | 3.989 | 0.00033 |
| males, 0% : $w^{-}$ vs. $FASN^{1\text{-}2}$ | 0.241 | 0.033 | 7.425 | 0.00000 |
| females, 0% : $w^{-}$ vs. $FASN^{1\text{-}2}$ | 0.263 | 0.032 | 8.257 | 0.00000 |
| males, 10% : $w^{-}$ vs. $FASN^{1\text{-}2}$ | 0.230 | 0.031 | 7.510 | 0.00000 |
| females, 10% : $w^{-}$ vs. $FASN^{1\text{-}2}$ | 0.272 | 0.031 | 8.884 | 0.00000 |

**Table S4. Statistical tests corresponding to Figure 6B.** Prepupal weight differences in males and females (Estimate and standard error, in mg) between control ($w^{-}$) and $FASN^{1\text{-}2}$ animals fed either a standard (0%) or a sucrose enriched diet (10%). The **Pr** column (right) indicates P-values corrected for multiple testing (Holm-Bonferoni correction).
